# Longitudinally persistent cerebrospinal fluid B-cells resist treatment in multiple sclerosis

**DOI:** 10.1101/490938

**Authors:** Ariele L. Greenfield, Ravi Dandekar, Akshaya Ramesh, Erica L. Eggers, Hao Wu, Sarah Laurent, William Harkin, Natalie S. Pierson, Martin S. Weber, Roland G. Henry, Antje Bischof, Bruce A.C. Cree, Stephen L. Hauser, Michael R. Wilson, H.-Christian von Büdingen

## Abstract

B-cells are key contributors to chronic autoimmune pathology in multiple sclerosis (MS). Clonally related B-cells exist in the cerebrospinal fluid (CSF), meninges, and central nervous system (CNS) parenchyma of MS patients. Longitudinally stable CSF oligoclonal band (OCB) antibody patterns suggest some local CNS B-cell persistence; however, the longitudinal B-cell dynamics within and between the CSF and blood remain unknown. We sought to address this by performing immunoglobulin heavy chain variable region repertoire sequencing on B-cells from longitudinally collected blood and CSF samples of MS patients (n=10). All patients were untreated at the time of the initial sampling; the majority (n=7) were treated with immune modulating therapies 1.2 (+/−0.3 SD) years later during the second sampling. We found clonal persistence of B-cells in the CSF of five patients; these B-cells were frequently immunoglobulin (Ig) class-switched and CD27+. We identified specific blood B-cell subsets that appear to provide input into CNS repertoires over time. We demonstrate complex patterns of clonal B-cell persistence in CSF and blood, even in patients on high-efficacy immune modulating therapy. Our findings support the concept that peripheral B-cell activation and CNS-compartmentalized immune mechanisms are in part therapy-resistant.

## Introduction

B lymphocytes (B-cells) are key players in the immunopathology of multiple sclerosis (MS) (1–6). B-cell depleting anti-CD20 antibody therapies effectively reduce relapsing MS disease activity and slow the accumulation of disability in relapsing and primary progressive disease *(2*, *3*, *7)*. Early on in the disease course, B-cells enter the CNS, CSF, and meningeal compartments (8–10) where they become compartmentalized and likely contribute to ongoing CNS tissue damage and consequential disability progression.

Immunoglobulin gene transcripts, particularly heavy chain variable regions (Ig-VH), provide molecular fingerprints that permit temporal and spatial tracking of clonally related B-cells (11–13). Single time-point immune repertoire studies in MS patients showed that there is an anatomic continuum of clonally related B-cells extending from MS lesions in the brain parenchyma, to cerebrospinal fluid (CSF), meningeal inflammatory infiltrates, cervical lymph nodes, and peripheral blood (PB) (11, 12, 14–16). Individual MS patients have a characteristic CSF immunoglobulin electrophoretic pattern (oligoclonal bands [OCBs]) that persists over many years; the stability in the pattern of OCBs within a patient is considered evidence that clonally related B-cells are long-lived in the central nervous system (CNS) (17, 18). However, because multiple B-cell clones can give rise to a single OCB (19), higher resolution genomic techniques are required to confirm this hypothesis.

Here, we were interested to determine persistence of clonally related B-cells in the CSF and in PB over time. Using previously described immune repertoire sequencing technology and bioinformatics tools (8), we analyzed Ig-VH sequences (IgG-VH and IgM-VH) (20) from 10 patients’ PB and CSF B-cell subsets at two time points to track related B-cells over time and across compartments. Work by others previously described two untreated MS patients with clonally related CSF B-cells over time (21). Here, we were interested to understand which specific B-cell subsets comprise clonally related CSF B-cells that remain longitudinally detectable, their clonal relationship to PB B-cells, and whether CSF B-cells persist after treatment with MS immune modulating therapy (IMT). We identified persistent clonally related B-cells in the CSF of five patients, four of whom had initiated IMT between the first and second time points. We also identified patterns of clonal B-cell input from the periphery to the CSF over time, suggesting that functionally diverse CD27+ PB memory B-cells are a likely peripheral reservoir of B-cells involved in MS disease activity.

## Results

### Patient demographics

10 patients ranging in age from 24 to 52 years old were enrolled (see Table 1 for clinical data and Table S1 for standard laboratory CSF findings). On average, 12.2 mL (+/− 3.5 SD) of CSF was collected per patient per time point. Eight patients had relapsing-remitting MS (RRMS), and two had primary progressive MS (PPMS). All were untreated at time point 1 (T1), and seven of the eight RRMS patients were on IMT by time point 2 (T2) (Table 1). Since there were no approved IMTs for PPMS prior to 2017, neither PPMS patient was on an IMT. IMTs used in this cohort varied from lower efficacy to higher efficacy treatments. Eight of 10 patients (6 treated, 2 untreated) had enhancing and/or new lesions on brain and/or spinal cord MRI at T2 (Table 1). All patients had OCBs unique to the CSF at both time points (Table S2). In 7/10 patients, we were able to directly compare the OCB pattern at each time point (Fig. S1); 5/7 patients had stable band patterns, one had a decrease in band number and one had an increase in band number (Fig. S1, Table S2).

**Table 1.**
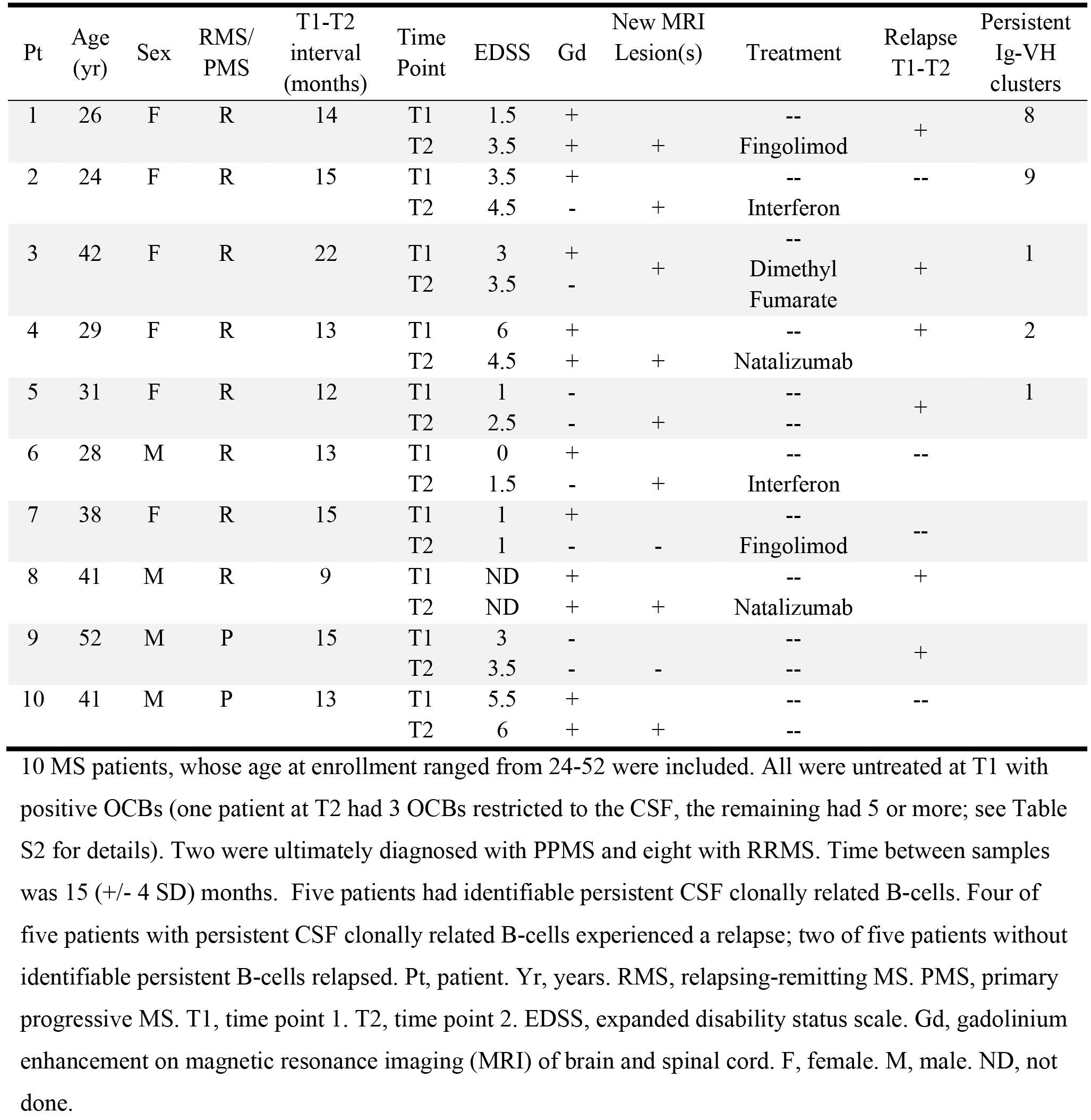
Patient Demographics

| Pt | Age (yr) | Sex | RMS/<br>PMS | T1-T2<br>interval<br>(months) | Time<br>Point | EDSS | Gd | New MRI<br>Lesion(s) | Treatment | Relapse<br>T1-T2 | Persistent<br>Ig-VH<br>clusters |
| --- | --- | --- | --- | --- | --- | --- | --- | --- | --- | --- | --- |
| 1 | 26 | F | R | 14 | T1 | 1.5 | + |  | -- | + | 8 |
|  |  |  |  |  | T2 | 3.5 | + | + | Fingolimod |  |  |
| 2 | 24 | F | R | 15 | T1 | 3.5 | + |  | -- | -- | 9 |
|  |  |  |  |  | T2 | 4.5 | - | + | Interferon |  |  |
| 3 | 42 | F | R | 22 | T1 | 3 | + | + | -- | + | 1 |
|  |  |  |  |  | T2 | 3.5 | - |  | Dimethyl Fumarate |  |  |
| 4 | 29 | F | R | 13 | T1 | 6 | + |  | -- | + | 2 |
|  |  |  |  |  | T2 | 4.5 | + | + | Natalizumab |  |  |
| 5 | 31 | F | R | 12 | T1 | 1 | - |  | -- | + | 1 |
|  |  |  |  |  | T2 | 2.5 | - | + | -- |  |  |
| 6 | 28 | M | R | 13 | T1 | 0 | + |  | -- | -- |  |
|  |  |  |  |  | T2 | 1.5 | - | + | Interferon |  |  |
| 7 | 38 | F | R | 15 | T1 | 1 | + |  | -- | -- |  |
|  |  |  |  |  | T2 | 1 | - | - | Fingolimod |  |  |
| 8 | 41 | M | R | 9 | T1 | ND | + |  | -- | + |  |
|  |  |  |  |  | T2 | ND | + | + | Natalizumab |  |  |
| 9 | 52 | M | P | 15 | T1 | 3 | - |  | -- | + |  |
|  |  |  |  |  | T2 | 3.5 | - | - | -- |  |  |
| 10 | 41 | M | P | 13 | T1 | 5.5 | + |  | -- | -- |  |
|  |  |  |  |  | T2 | 6 | + | + | -- |  |  |

### Flow Cytometry

B-cell subsets were identified and sorted by flow cytometry (MoFlo Astrios). Flow cytometric B-cell subset distribution (see Methods) was determined at T1 and T2 for n=8 patients’ CSF and for n=9 patients’ PB; flow cytometry was not available for time point 1 CSF (T1-CSF) and peripheral blood (T1-PB) of one patient and for both CSF time points of another patient. When compared to PB, the CSF was enriched in CD19+CD27+IgD-Ig class-switched memory (SM) (Fig. S2), consistent with previous reports (8, 22, 23).

### Immune Repertoire Sequencing

IgG-VH and/or IgM-VH repertoire sequencing (IgSeq) cDNA libraries were prepared from 167 samples. Samples consisted of PB or CSF sorted B-cell subsets or alternatively bulk CSF or PB mononuclear cells (Table S3). Sequencing libraries could not be obtained from 16 samples (Table S3). From the remaining 151 samples, we generated 583,932 (+/− 652,920 SD) raw reads per library. 218,401 (+/− 308,602 SD) Ig-VH sequences per library were identified from the immunoglobulin heavy chain variable germline segment (IGHV), immunoglobulin heavy chain joining germline segment (IGHJ) and Ig heavy chain complementarity-determining region 3 (H-CDR3) for further analysis (Table S3). Ig-VH sequences were clustered using a distance metric approach (see Methods). Samples with more B-cells had more Ig-VH clusters (Fig. S3) (r = 0.88, p<0.0001 for all samples; r = 0.74, p<0.0001 for PB; r = 0.59, p<0.0001 for CSF, Spearman correlation). Five paucicellular B-cell subsets yielded more Ig-VH clusters than the number of input cells (Table S3). For these samples, we analyzed the same number of Ig-VH clusters as input cells, choosing the Ig-VH clusters with the greatest number of aligned sequencing reads. Mutational analyses within Ig-VH clusters were not performed as these were not needed for the conclusions of this study.

At T1, we identified CSF Ig-VH clusters that were exclusively IgG-VH in all 10 patients (26.4 (+/− 28.3 SD) Ig-VH clusters/patient); of the 10 patients, nine patients also had CSF Ig-VH clusters that contained exclusively IgM-VH (44.8 (+/− 57.3 SD) Ig-VH clusters/patient) and in five patients, we found mixed IgM and IgG clusters 5.2 (+/− 9.4 SD) Ig-VH clusters/patient (Fig. S4). At T2, we found that all 10 patients’ CSF contained Ig-VH clusters that were exclusively IgG-VH (42.6 (+/− 72.6 SD) Ig-VH clusters/patient) or exclusively IgM-VH (31.7 (+/− 33.9 SD) Ig-VH clusters/patient); in seven patients, there were 6.7 (+/− 11.8 SD) Ig-VH clusters/patient that were mixed IgM and IgG (Fig. S4). At T1, SM and naïve B-cells were common members of CSF Ig-VH repertoire clusters: these subsets were prevalent in repertories of 3/5 and 2/5 patients with sorted CSF B-cells at T1 (T1-CSF) B-cells, respectively (59.6 +/− 80.4 SD, 60.8 +/− 65.9 SD Ig-VH clusters/patient, respectively; Fig. S5). SM B-cells commonly contributed to T2-CSF: 5/8 patients with sorted T2-CSF B-cells had SM-predominant repertoires (45.4 +/− 65.9 SD Ig-VH clusters/patient) (Fig. S5).

### Clonally related B-cells persist in MS CSF

In 5 of 10 patients, we identified ‘persistent CSF Ig-VH clusters’ in which CSF Ig-VH sequences from both time points were represented (Figs. 1-3); we thus demonstrate that B-cells found in MS CSF at different time points are clonally related. Aside from Ig-VH sequences that exclusively persisted in CSF (Fig. S6), we identified three possible associations of CSF Ig-VH clusters with PB repertoires: a T1-PB connection, T2-PB connection, or connections with both PB time point samples (Fig. 2). We found IgG-expressing B-cells, including SM and PC, in persistent CSF Ig-VH clusters of all 5 patients with persistent CSF Ig-VH clusters (Fig. 3). In contrast, IgM-expressing B-cell subsets were only found to take part in persistent CSF Ig-VH clusters in two patients (patients 1 and 3) (Fig. 3). In particular, we did not find naïve CSF B-cells in persistent CSF Ig-VH clusters.

**Figure 1.**
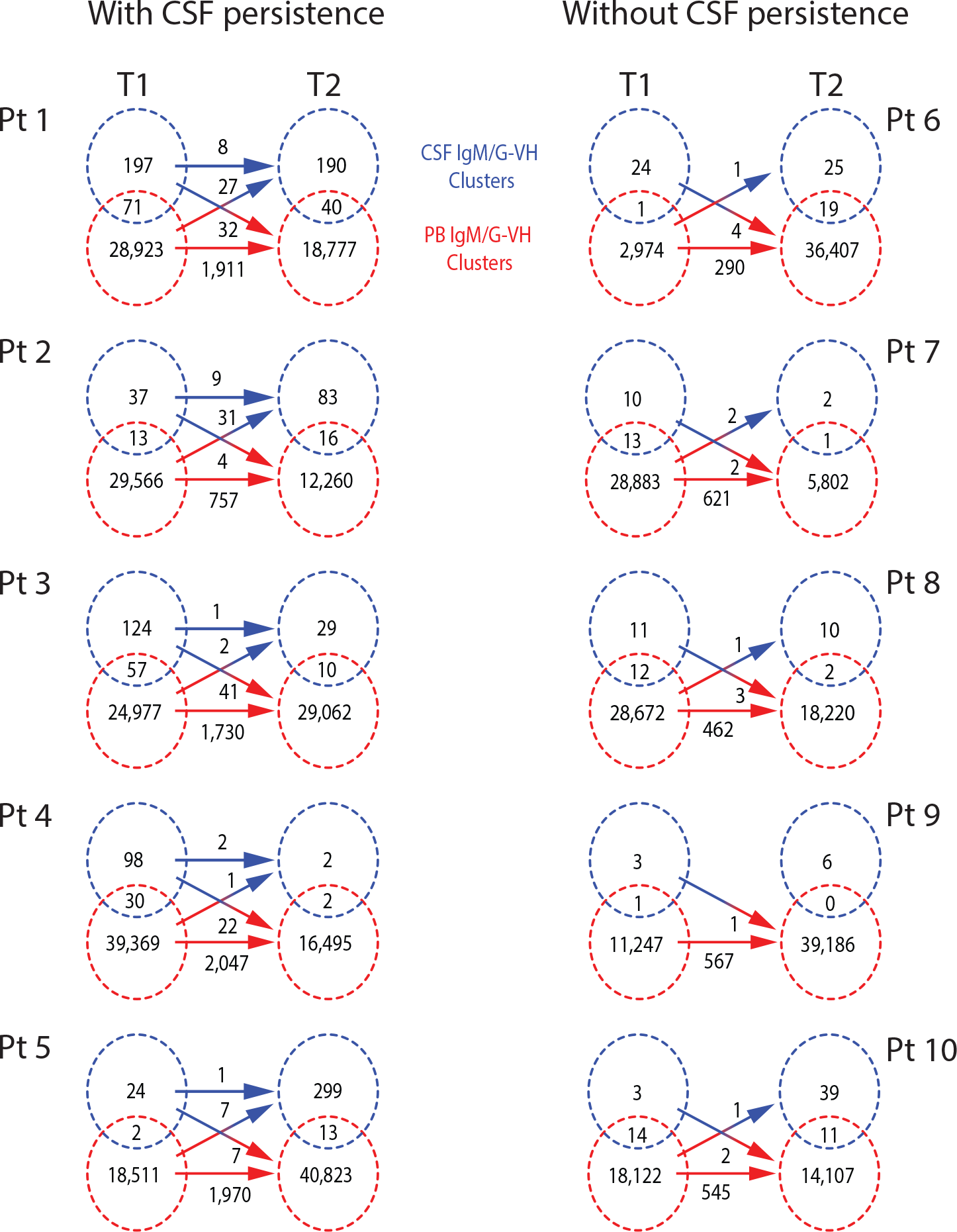
Persistent CSF Ig-VH clusters are present in MS patients. Shown are the numbers of Ig-VH clusters within CSF (blue circles) and PB (red circles) at T1 and T2. Circle overlap values - Ig-VH clusters found in both CSF and PB at T1 or T2; non-overlap circle values - Ig-VH clusters exclusively found in CSF or PB, not both. Arrows - of total Ig-VH clusters in CSF or PB (non-overlap portion of circle + circle overlap), the number of Ig-VH clusters that are present at both T1 and T2. Pts 1-5 have persistent CSF Ig-VH clusters; Pts 6-10 did not have identifiable persistent CSF Ig-VH clusters. Pt, patient. Ig-VH, immunoglobulin heavy chain variable region. IgG, immunoglobulin G. IgM, immunoglobulin M. CSF, cerebrospinal fluid. PB, peripheral blood.

**Figure 2.**
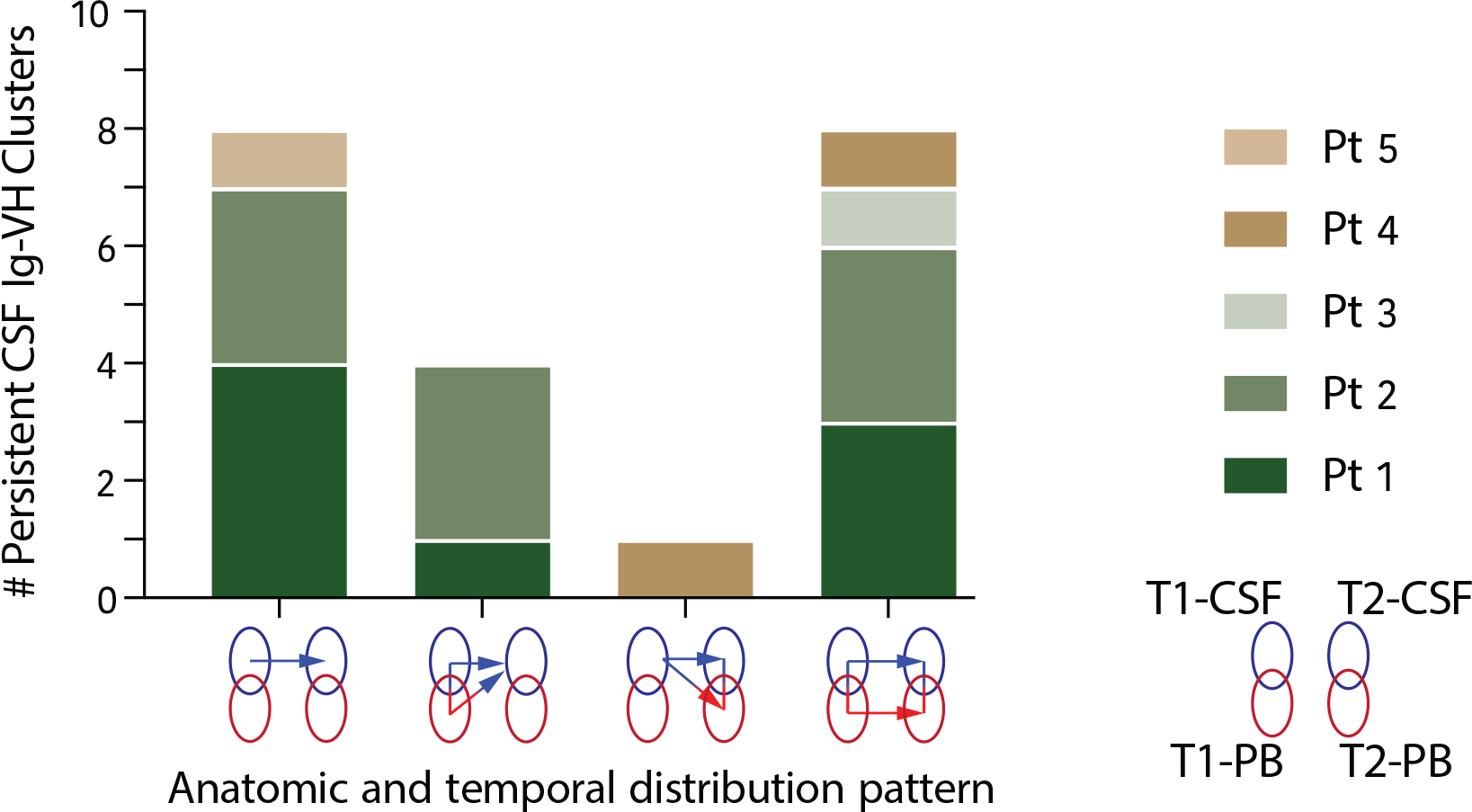
Persistent CSF Ig-VH clusters have multiple patterns of persistence in relation to PB. Persistent CSF Ig-VH clusters can be exclusive to CSF (far left: Patients 1,2,5), link to T1-PB (second from left: Pts 1,2), link to T2-PB (second from right: Patient 4), or link to both PB time points (far right: Pts 1-4). Blue circles: CSF. Red circles: PB. Circles on left: T1. Circles on right: T2. Arrows: shared Ig-VH clusters between time points or between CSF and PB.

**Figure 3.**
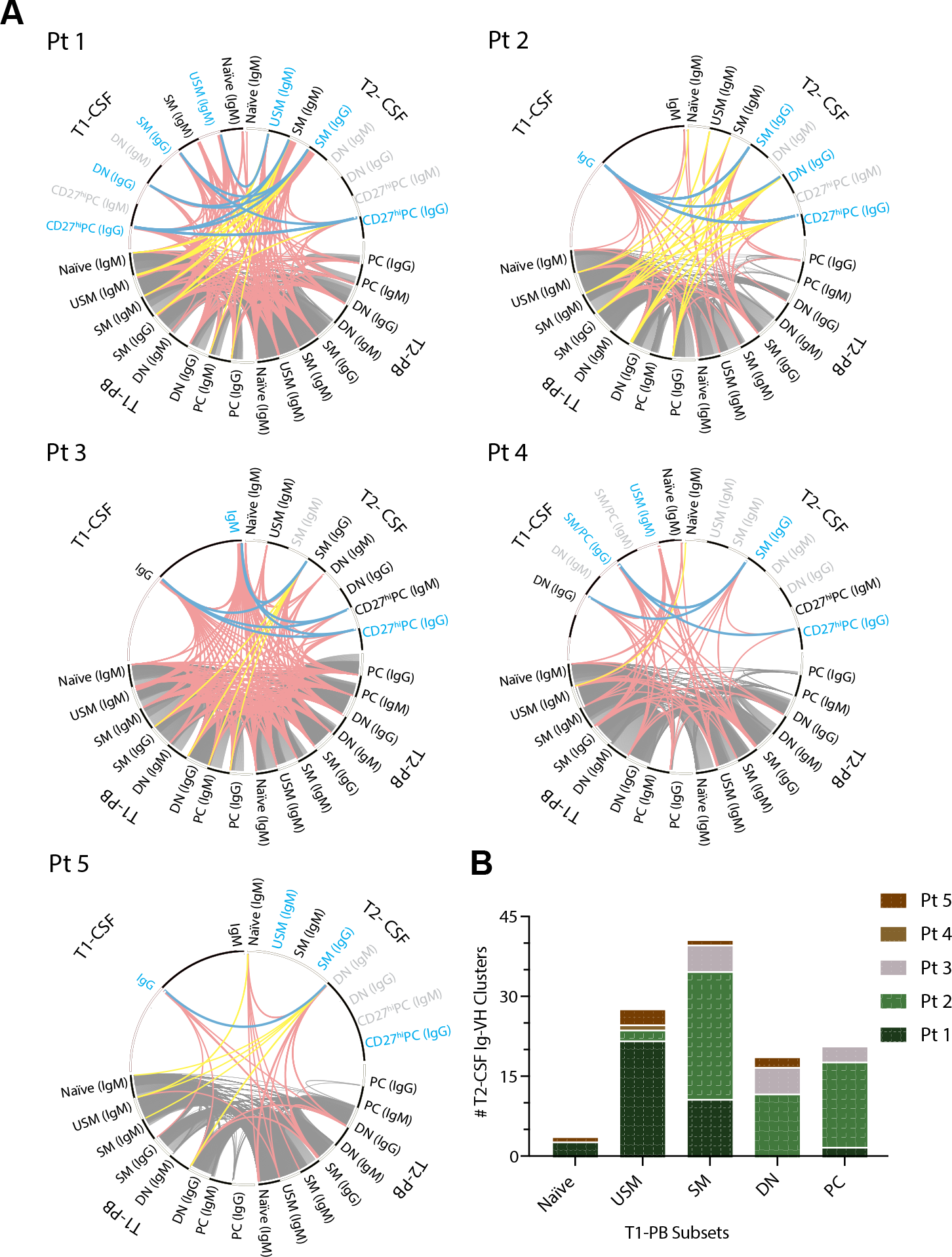
B-cells that persist in the CSF are often SM and PC. A) Clonally-related B-cell subsets are connected by lines between CSF (top half of circle) and PB (bottom half) at T1 (left half) and T2 (right half). Each segment along the circle represents a B-cell subset from CSF or PB. Lines connecting B-cell subsets indicate B-cell subsets that share one or more Ig-VH clusters in common. Grey lines: PB-only Ig-VH clusters. Blue lines: Persistent CSF Ig-VH clusters. Red lines: CSF Ig-VH clusters with PB contribution. Yellow lines: T1-PB Ig-VH clusters that share clonal relatives with T2-CSF yet not with T1-CSF. Grey font indicates subsets/Ig isotypes from which no Ig-VH libraries could be obtained. Persistent CSF Ig-VH clusters were linked to PB Ig-VH sequences derived from SM (n=4 patients), DN (n=3 patients at T1, n=2 patients at T2), and PC (n=2 patients at T1 and 1 patient at T2). B) T1-PB B-cell subsets that are clonally related to T2-CSF without involving T1-CSF (i.e. quantitation of yellow lines depicted in (A)). USM, unswitched memory. SM, class-switched memory. DN, double negative. PC, plasmablast/plasma cell.

There was no relationship between the total CSF white blood cell count (clinical diagnostic which includes all leukocytes, not just B-cells) at T1 or T2 and the number of persistent CSF Ig-VH clusters (r^2^=0.32, p=0.09 and r^2^;0.07, p<0.47, respectively; not shown). There was no significant correlation between the detection of persistent CSF Ig-VH clusters over time and the following clinical characteristics: age, disease duration, time between T1 and T2, expanded disability status scale (EDSS), use of IMT, gadolinium-enhancement on MRI at either time point, IgG index, relapse between T1 and T2, or number of OCBs (Table S1, Table S2). More of the patients without persistent CSF Ig-VH clusters were men. There was a numerical (yet not statistically significant) imbalance in the number of patients with relapses in that more of the patients with identified persistent CSF Ig-VH clusters experienced a relapse. There was no significant difference between B-cell subset distribution in patients with or without persistent CSF Ig-VH clusters (Fig. S7). We found no correlation between the average reads per cell and our ability to detect persistent CSF Ig-VH clusters (r=0.2, Spearman correlation; p=0.17). The percentage of Ig-VH clusters in CSF that were persistent was similar to the percentage of Ig-VH clusters that were persistent in PB (i.e. Ig-VH clusters formed from PB at both time points) (Table S4). This suggests that immune processes such as antigen stimulation and B-cell regulation, which maintain a proportion of B-cell clonal populations in the periphery, may also occur in the CNS.

### PB SM and Unswitched Memory (USM) B-cells - potential contributors to CSF B-cell repertoires

Clonal connections between PB and CSF were identified in all patients, irrespective of whether persistent CSF Ig-VH clusters, or clonal relatedness within the same PB and CSF subset, was present (Figs. 3 and S8). Therefore, to understand which PB B-cell subsets might contribute clonal input to CSF repertoires over time we examined Ig-VH clusters containing T1-PB and T2-CSF members, yet no T1-CSF members. In all 9 patients with sorted T1-PB subsets, T1-PB USM (6 patients) and SM (5 patients) B-cells were related to T2-CSF (Fig. 3, Fig. S8). All of these patients except for one (Pt 5) were treated with IMT by T2. Other T1-PB B-cell subsets sharing repertoires with T2 CSF included naïve (4 patients), DN (3 patients), and PC (3 patients) (Fig. 3, Fig. S8). While overall somatic hypermutation (SHM) rates followed expected patterns for B-cell subsets based on maturation stage, certain CSF subsets, such as IgG SM in Patient 1, were noted to have a particularly high degree of SHM - suggesting affinity maturation (Fig. S9). Whether Ig-class switching occurs in USM B-cells has been debated (24); here we found numerous Ig-VH clusters containing both USM (CD27+IgD+IgM+) and class-switched IgG+ B-cells (i.e. SM, PC, double negative [DN]) in CSF and PB (2.5 +/− 2.6 SD Ig-VH clusters/patient in CSF, 125.6 +/− 96.6 SD Ig-VH clusters/patient in PB) (Fig. S10). To explore whether CSF PCs might be derived from long-lived PB PC populations, we searched for, but did not find, persistent PB PCs that were clonally related to CSF PC. Incidentally, we found frequent occurrence of persisting IgM-VH clusters with PB naïve B-cell members at T1 and T2, suggesting long-term survival and clonal expansion of naïve B-cells; previously, naïve B-cells have been characterized as short-lived precursors to memory and plasmablast/plasma cells (25).

## Discussion

We identified clonally related B-cells in CSF that were detectable at time-points between 9 and 22 months apart. Given the histopathological and biochemical evidence for B-cell recruitment, survival, and proliferation in the CNS compartment (26) (27) (28), the ability to detect clonally related CSF B-cells by immune repertoire sequencing over time is likely the result of local persistence (i.e. long-term survival) of B-cells in the functionally connected CNS, CSF, and meningeal compartments. These cells could constitute a tissue-resident immune repertoire capable of being re-stimulated - either directly or by bystander activation - to re-kindle inflammation; mechanisms that may be shared with rheumatoid arthritis and other tissue-specific chronic autoimmune diseases (29). B-cells of persistent CSF Ig-VH clusters display phenotypic features indicative of antigen-driven immunity, i.e. CD27-expression and Ig class-switched B-cell receptors. Persistent B-cells in PB, on the other hand, belong to a less restricted set of B-cell subsets and include apparently long-lived naïve (IgD+CD27-) B-cell populations in addition to memory B-cell and PC subsets. Overall, persistent CSF B-cells comprise a minority of the overall CSF B-cell diversity.

Nonetheless, their persistence despite immune modulatory treatment initiation suggests that these B-cell populations could be representative of a therapy-resistant CNS compartmentalized reservoir of immune cells. OCBs as well are evidence for CNS/CSF compartmentalization, albeit only of plasma cells as producers of soluble immunoglobulin. CNS compartmentalized B-cells are believed to contribute to progressive CNS tissue damage in MS (26, 30, 31); identification of CSF persistent functionally diverse B-cell subsets (e.g. memory B-cells, plasma cells) suggests that multiple aspects of active adaptive immunity are reflected in the CNS capable of supporting the immunopathogenesis of MS in various ways (i.e. cytokine secretion, antigen presentation, and antibody production). Interestingly, even after long-term high efficacy IMT (32, 33), including intrathecal rituximab (34), patients continue to be at risk of progression. Indeed, our findings lend further support to the hypothesis that currently available IMTs may not sufficiently target progression-promoting mechanisms located in the CNS compartment. Future studies are needed to determine whether anti-CD20 therapies also deplete persistent clonal B-cells from the CSF, CNS, and meninges and whether doing so reduces disease progression. Our findings may suggest that development of novel therapies that target CNS compartmentalized immune mechanisms could help close an important therapeutic gap.

Disease-driving lymphocytes, including B-cells, are believed to migrate to the CSF and CNS from peripheral stores during phases of MS disease activity (8, 11, 12). Due to their immunological properties and functionality, SM B-cells are generally believed to be highly relevant to the immune pathology of MS (35). Indeed, when persistent PB SM cells were clonally related to second time-point CSF B-cells, these T2-CSF cells were nearly always SM and PC. This suggests that, in addition to CSF persistence, a population of potentially disease-relevant PB SM B-cells may have a particular tendency to migrate to the CSF and mature to PC. Also suggestive of peripheral maturation to SM followed by migration and survival in the CNS, rather than local SM to PC maturation in CSF/CNS, we did not find any T2-CSF B-cells that were clonally related to T1-CSF naïve B-cells. In agreement with our previous work, we did not observe any relationships between PB-persistent PCs and CSF PC, further supporting that migration to the CNS occurs at the SM, or earlier B-cell development stage followed by intrathecal maturation to PC rather than immigration of long-lived peripheral PCs.

The efficacy of anti-CD20 B-cell depleting therapy in autoimmune disease is linked to depletion of memory B-cell subsets rather than naïve and transitional subsets (36, 37). We previously proposed that USM B-cells may play a role in MS due to their appearance in MS CSF during periods of active CNS inflammation (8). Interestingly, USM B-cells follow the same depletion and reconstitution pattern as SM B-cells under rituximab therapy (38). Here, we observed that clonally related SM and USM B-cells, which remain detectable in PB over time, may also have clonal relatives in CSF at the later time point. This finding is consistent with the concept that, upon peripheral activation, CXCR5+ USM and/or SM B-cells home to the brain parenchyma or leptomeninges along a CXCL13 gradient, where they participate in the formation of ectopic, tertiary germinal centers (8, 31, 39, 40) capable of supporting B-cell receptor affinity-maturation and class-switch recombination (41). Whether USM B-cell receptors undergo Ig class-switching and contribute to high-affinity surface receptor or secreted antibody-mediated immune responses has been debated (42, 43). Here, we found numerous Ig-VH clusters linking CSF or PB IgM-expressing USM to clonally related IgG+ B-cells in CSF and PB (i.e. SM, PC, DN). This finding suggests either 1) that USM B-cells can enter germinal centers where they undergo class-switch recombination, or 2) that divergent maturation occurs of naïve B-cells into functionally diverse IgM- and IgG-expressing memory B-cell subsets. Similar to SM B-cells (44), USM B-cells can likely function as potent antigen-presenting cells (45). Therefore, the depletion of both SM and USM B-cells could be important for the efficacy of B-cell depleting MS therapies. However, whether USM B-cells are involved in a true antigen-directed immune response or, conversely, whether they provide a non-specific stimulus for intrathecal immune activation remains unclear. In either case, due to their participation in clonally-related immune connectivity between PB and CSF, SM and USM B-cells may be carriers of MS disease activity.

There are a number of limitations to this study. First, it is important to keep in mind that it is not technically possible to interrogate the entire immune repertoire in a living patient. As a result, persistent B-cells could have gone undetected, and patients in whom we did not find clusters comprised of clonally related Ig-VH may in fact harbor persistent B-cells in the CSF, or in other tissues. While it is possible that deeper sequencing of existing samples could have increased our sensitivity to identify persistent CSF Ig-VH clusters, and while there were some samples that did not generate productive sequences, we found no correlation between the average reads per cell and our ability to detect these clusters. Second, technical advances such as incorporation of unique molecular identifiers (UMIs) that aid in error correction and quantification were not a standard part of the sequencing method at the start of the study, and the initial method was maintained for consistency across this cohort. PCR or sequencing errors are unlikely to influence determination of presence versus absence of related Ig-VH sequences, but may influence the quantity of Ig-VH clusters. For this reason, we chose a distance metric (see Methods) that minimizes risk of overestimating the number of Ig-VH clusters that could result from sequencing errors introduced by high-throughput technology. Third, although to our knowledge, this is the largest study of sorted B-cell subsets from paired CSF and PB samples evaluated at two time points, only 10 patients were studied. Future studies will be needed to confirm these findings in larger cohorts, including patients treated with B-cell depleting therapies. Fourth, our experimental approach does not allow us to determine the directionality of B-cell migration; to what extent lymphocyte migration pathways via ‘glymphatics’ (46, 47) underlie our findings remains subject to future studies that will require reliable *in vivo* lymphocyte tracking techniques. The question whether clonally related B-cells mature and persist in the CNS or mature and then migrate from the periphery is an important one. However, we refrain from performing mutational lineage analyses to determine B-cell migration directionality, as intermediate mutants that are necessary to obtain a complete picture of Ig-VH evolution may have gone undetected in our study.

In summary, we describe CSF persistent clonally related B-cell populations as part of an overall complex picture of B-cell relatedness over time and across compartments that appears at least partially independent of IMT initiation. We identified previously unknown patterns of B-cell relatedness, including PB SM and USM B-cells that may contribute to SM- and PC-rich CSF B-cell repertoires, and naïve B-cells that persist over time. Our findings raise the possibility that IMTs which effectively reduce clinical and MRI relapses may not sufficiently target CNS compartmentalized immune mechanisms that have been associated with gradual and relapse-independent MS worsening in relapsing MS and progressive forms of the disease (48). Effective targeting of CNS compartmentalized inflammation may, in the future, achieve efficacy on progression slowing superior to that of current therapies (2). Longitudinal immune repertoire studies performed alongside the clinical development of such novel MS drugs may well provide critical immunologic and biomarker evidence indicating the successful suppression or elimination of progression-driving mechanisms.

## Materials and Methods

### Experimental Design

We hypothesized prior to initiating this study that in MS patients there would be a population of B-cell clones that persist in the CSF over time. Inclusion criteria for this observational study were 1) enrollment in the University of California at San Francisco (UCSF) Expression, Proteomics, Imaging, Clinical (EPIC) Study (33) which is a longitudinal MS cohort that defines MS according to the 2001 McDonald criteria (49), and 2) biobanked CSF and PB mononuclear cells (PBMCs) from two time points (T1 and T2) taken a minimum of 9 months apart. All patients were treatment-naïve at T1. Patients either consented to donating excess CSF and blood during routine lumbar puncture (LP) at UCSF; if an LP was not ordered by the treating physician, participants consented to undergo LP and blood draw for research purposes. Patient demographics, clinical history, including physical exam and EDSS scores, were also obtained as part of the EPIC study.

### Sample Collection and Processing

7-30 mL of fresh CSF was centrifuged at 400g × 15 minutes at 4°C to separate a cell pellet from supernatant. PBMCs were isolated from whole blood via Ficoll gradient followed by red blood cell lysis and washing with phosphate buffered saline with 1% bovine serum albumin. Seven of 20 CSF samples and one of 20 PBMC samples were stored at -80°C as unsorted cell pellets (Table S3). The remaining CSF and PBMC samples were immediately blocked with FcR Block (Miltenyi Biotec, Bergisch Gladbach, Germany) and stained with fluorescent antibodies to cell surface markers - CD19, CD27, CD38, CD138, IgD; some samples were also stained for additional immune cell markers for experiments outside the scope of this study (antibody panels listed in Table S5, Table S6).

CSF and PB B-cell subsets were sorted as previously described (8) on a Beckman MoFlo Astrios fluorescent-activated cell sorting (FACS) sorter: naïve B-cells (N: CD19+IgD+CD27-), unswitched memory B-cells (USM: CD19+IgD+CD27+), class-switched memory B-cells (SM: CD19+IgD-CD27+), double negative B-cells (DN: CD19+IgD-CD27-), CSF CD27^hi^ plasmablast/plasma cells (PC: CD19+IgD-CD27^hi^), the vast majority of which are also CD38+, and PB plasmablast/plasma cells (PC: CD19+IgD-CD27+CD38+). Patient 4 had combined CD^27+/hi^ CSF sorted B-cells. CSF samples with concern for low B-cell count were bulk sorted - attaining flow cytometry data, but placing all CSF lymphocytes into a single sample tube rather than segregating into the above five subsets. B-cells were sorted directly into lysis buffer suitable for later RNA extraction (Qiagen, buffer RLT+ 1% β-mercaptoethanol) and stored at -80°C.

### Immune Repertoire Sequencing

RNA was extracted using the RNeasy Mini kit for >200,000 cells or the Micro kit for <200,000 cells (Qiagen, Hilden, Germany). RNA was reverse transcribed using iScript reverse transcriptase (BioRad, Hercules, CA). IgG-VH and IgM-VH sequences spanning the variable region from framework 1 (IGHV FR1) to the constant region 1 were amplified using the Advantage2 PCR kit (Clontech, Mountain View, CA) and custom primers as described (8). 30 PCR cycles were performed on PB samples and 40 cycles on CSF samples; up to 10 additional cycles were performed, in 5 cycle increments, if no PCR product was visualized on a 1.5% agarose gel after the initial round of amplification. Amplicons of the expected length were gel purified and quantified by Bioanalyzer using the High Sensitivity DNA kit (Agilent, Santa Clara, CA). Equimolar amounts of PCR products were pooled to generate a 16pM library for IonTorrent sequencing (Thermo Fisher, Waltham, MA) following the manufacturer’s instructions, including emulsion PCR (Ion OneTouch2), enrichment for template-positive ion sphere particles, and quality control using a Qubit fluorometer (IonTorrent QC kit). Next-generation sequencing was performed on an IonTorrent Personal Genome Machine. If inadequate sequencing data was obtained during an initial sample’s sequencing, a technical replicate was performed.

### Bioinformatics

Raw sequence FASTQ files were generated with Torrent Suite software (Thermo Fisher, Waltham, MA). A custom bioinformatics pipeline incorporating MiXCR (v2.1.3) (50) was used to identify IGHV and IGHJ germline segments and H-CDR3 for each sequence read. MiXCR first assembles reads with bases of quality scores >20 and then attempts to align reads with bases of lower (<20) quality to the already assembled reads. We applied a bioinformatics clustering approach as described previously (8) to compile clonally related Ig-VH sequences into ‘Ig-VH clusters’. An Ig-VH cluster contains clonally related sequences - IgG and/or IgM - from at least one B-cell. B-cell receptors using identical IGHV and IGHJ germline segments and identical or near-identical H-CDR3 regions (minimum 8 amino acid length, maximum Hamming distance (51) of 2 at the amino acid level were considered members of a clonally related Ig-VH cluster. This distance metric permitted clustering of identical and clonally related Ig-VH irrespective of sequencing errors that might have introduced a frameshift in the Ig-VH. Immune repertoire datasets can have falsely inflated diversity as a result of errors during the reverse transcription, amplification and sequencing steps as well as incorrect barcode assignment that occurs when sequencing libraries are pooled. Thus, to conservatively catalog immune repertoire diversity in a given sample, we excluded any clusters that had fewer than 2 aligned read counts and any additional, low abundance Ig-VH clusters whose inclusion resulted in the total number of clusters exceeding the number of input cells. For samples in which library generation was unsuccessful, we excluded these samples from statistical analyses.

Network diagrams were generated to visualize Ig-VH clusters using Circos (52). Amino acid alignment figures were generated using Geneious 10.0.6 (https://www.geneious.com) and IMGT High-V Quest (53). Venn diagrams and other figures were based on custom R scripts that tabulated the number and type of Ig-VH clusters shared across compartments (CSF/PB) and time points.

### MRI Acquisition and Analysis

MRI scans at T1 and T2 were acquired on the same 3T Siemens scanner following a standardized protocol that included: 3D T1-weighted magnetization-prepared gradient echo images (MPRAGE), 3D fluid-attenuated inversion recovery (FLAIR) and T2/PD images of the brain; whole spine T2-weighted and short tau inversion recovery (STIR) images. In addition, brain MPRAGE and whole spine T1-weighted images were acquired after administration of gadobutrol. Readings were performed by a neurologist with subspecialty interest in neuroimaging (AB, 11 years of experience) blinded to the clinical and immunological data.

### Statistical Analysis

Graphpad Prism software (version 8) was used to perform statistical analyses: two-tailed unpaired t-tests were used to compare pairwise groups and Fisher’s exact test was used to compare categorical variables. ANOVA was used for comparisons between groups with approximate Gaussian distributions, with correction for multiple comparisons performed using the Sidak method. Kruskal-Wallis test with Dunn correction for multiple comparisons was used for multiple comparisons with non-parametric distributions. Spearman correlation coefficient was used to determine correlation. p<0.05 was considered statistically significant.

### Informed Consent

All studies were approved by the UCSF Institutional Review Board, and written informed consent was obtained from each participant prior to inclusion in the study.

## Supporting information

## Acknowledgments

The authors would like to express their gratitude to the individuals who agreed to participate as patients in this study, as well as the UCSF MS Expression, Proteomics, Imaging, Clinical (EPIC) study team of researchers and clinicians.

## Author contributions

ALG performed immune repertoire sequencing, analysis and writing of the manuscript; HCvB conceptualized the study, assisted in patient recruitment, provided guidance on analysis, and wrote the manuscript; MRW wrote the manuscript and provided guidance on analysis; RD and HW generated code, tables and figures for analysis; EE performed FACS and immune repertoire sequencing; MSW performed the OCB analysis; AB performed the analyses of MRI data; RGH oversaw acquisition of MRI data; SL performed immune repertoire sequencing and input on analysis; WH and NP assisted in patient recruitment, consenting and sample collection; BAC led the EPIC cohort study; SLH conceived of and led the EPIC cohort study.

## Data and materials availability

Immunoglobulin sequencing data will be deposited in the NCBI Sequence Read Archive (Bioproject accession number PRJNA397295).

## Supplementary Materials

Figure S1. CSF-unique OCBs are mostly stable over time.

Figure S2: naïve B-cells are more prevalent in blood; in CSF, SM B-cells are relatively increased.

Figure S3. Number of Ig-VH clusters in a sample is correlated with cell count.

Figure S4. The majority of CSF immune repertoire Ig-VH clusters express either IgM or IgG

Figure S5. Different B-cell subsets comprise CSF Ig-VH repertoires at T1 and T2

Figure S6. Amino acid alignment of a CSF-persistent Ig-VH cluster

Figure S7. Patients with persistent CSF Ig-VH clusters show no significant difference in B-cell type prevalence in CSF or PB compared to patients without persistent CSF Ig-VH clusters.

Figure S8. Patients without identifiable persistent CSF Ig-VH clusters still have clonal connections between CSF and PB.

Figure S9. Somatic hypermutation rates follow expected patterns along B-cell lineage.

Figure S10: Clonal relationships between IgM-expressing USM B-cells and IgG-expressing B-cell subsets suggest Ig class-switch recombination and further maturation of USM B-cells.

Table S1. Patient characteristics based on presence or absence of CSF persistent Ig-VH

Table S2. Clinical CSF biometrics

Table S3. B-cell samples analyzed by IgSeq

Table S4: CSF Ig-VH cluster persistence rate is similar to PB Ig-VH cluster persistence rate.

Table S5. FACS antibody sort panels

Table S6. FACS antibody sort panels used on cerebrospinal fluid and peripheral blood

## References

1. Molnarfi N, Schulze-Topphoff U, Weber MS, Patarroyo JC, Prod’homme T, Varrin-Doyer M, et al. MHC class II-dependent B cell APC function is required for induction of CNS autoimmunity independent of myelin-specific antibodies. J Exp Med. 2013;210(13):2921–37.

2. Montalban X, Hauser SL, Kappos L, Arnold DL, Bar-Or A, Comi G, et al. Ocrelizumab versus Placebo in Primary Progressive Multiple Sclerosis. N Engl J Med. 2017;376(3):209–20.

3. Hauser SL, Bar-Or A, Comi G, Giovannoni G, Hartung HP, Hemmer B, et al. Ocrelizumab versus Interferon Beta-1a in Relapsing Multiple Sclerosis. N Engl J Med. 2017;376(3):221–34.

4. Hauser SL. The Charcot Lecture | beating MS: a story of B cells, with twists and turns. Multiple sclerosis (Houndmills, Basingstoke, England). 2015;21(1):8–21.

5. Dalakas MC. B cells as therapeutic targets in autoimmune neurological disorders. Nature clinical practice Neurology. 2008;4(10):557–67.

6. t Hart BA, van Meurs M, Brok HP, Massacesi L, Bauer J, Boon L, et al. A new primate model for multiple sclerosis in the common marmoset. Immunology today. 2000;21(6):290–7.

7. Hawker K, O’Connor P, Freedman MS, Calabresi PA, Antel J, Simon J, et al. Rituximab in patients with primary progressive multiple sclerosis: results of a randomized double-blind placebo-controlled multicenter trial. Annals of neurology. 2009;66(4):460–71.

8. Eggers EL, Michel BA, Wu H, Wang S-z, Bevan CJ, Abounasr A, et al. Clonal relationships of CSF B cells in treatment-naïve multiple sclerosis patients. JCII nsight. 2017;2(22).

9. Frischer JM, Bramow S, Dal-Bianco A, Lucchinetti CF, Rauschka H, Schmidbauer M, et al. The relation between inflammation and neurodegeneration in multiple sclerosis brains. Brain : a journal of neurology. 2009;132(Pt 5):1175–89.

10. Bevan RJ, Evans R, Griffiths L, Watkins LM, Rees MI, Magliozzi R, et al. MENINGEAL INFLAMMATION AND CORTICAL DEMYELINATION IN ACUTE MULTIPLE SCLEROSIS. Annals of neurology. 0(ja).

11. Stern JN, Yaari G, Vander Heiden JA, Church G, Donahue WF, Hintzen RQ, et al. B cells populating the multiple sclerosis brain mature in the draining cervical lymph nodes. Science translational medicine. 2014;6(248): 248ra107.

12. Palanichamy A, Apeltsin L, Kuo TC, Sirota M, Wang S, Pitts SJ, et al. Immunoglobulin class-switched B cells form an active immune axis between CNS and periphery in multiple sclerosis. Science translational medicine. 2014;6(248):248ra106.

13. Bashford-Rogers RJM, Nicolaou KA, Bartram J, Goulden NJ, Loizou L, Koumas L, et al. Eye on the B-ALL: B-cell receptor repertoires reveal persistence of numerous B-lymphoblastic leukemia subclones from diagnosis to relapse. Leukemia. 2016;30(12):2312–21.

14. von Budingen HC, Gulati M, Kuenzle S, Fischer K, Rupprecht TA, and Goebels N. Clonally expanded plasma cells in the cerebrospinal fluid of patients with central nervous system autoimmune demyelination produce “oligoclonal bands”. Journal of neuroimmunology. 2010;218(1-2): 134–9.

15. Obermeier B, Lovato L, Mentele R, Bruck W, Forne I, Imhof A, et al. Related B cell clones that populate the CSF and CNS of patients with multiple sclerosis produce CSF immunoglobulin. J Neuroimmunol. 2011;233(1-2):245–8.

16. Bankoti J, Apeltsin L, Hauser SL, Allen S, Albertolle ME, Witkowska HE, et al. In multiple sclerosis, oligoclonal bands connect to peripheral B-cell responses. Annals of neurology. 2014;75(2):266–76.

17. Yu X, Burgoon M, Green M, Barmina O, Dennison K, Pointon T, et al. Intrathecally synthesized IgG in multiple sclerosis cerebrospinal fluid recognizes identical epitopes over time. J Neuroimmunol. 2011;240-241:129-36.

18. Walsh MJ, and Tourtellotte WW. Temporal invariance and clonal uniformity of brain and cerebrospinal IgG, IgA, and IgM in multiple sclerosis. J Exp Med. 1986; 163(1):41–53.

19. Bankoti J, Apeltsin L, Hauser SL, Allen S, Albertolle ME, Witkowska HE, et al. In multiple sclerosis, oligoclonal bands connect to peripheral B-cell responses. Annals of neurology. 2014;75(2):266–76.

20. Glanville J, Kuo TC, von Büdingen H-C, Guey L, Berka J, Sundar PD, et al. naïve antibody gene-segment frequencies are heritable and unaltered by chronic lymphocyte ablation. Proceedings of the National Academy of Sciences. 2011;108(50):20066–71.

21. Colombo M, Dono M, Gazzola P, Chiorazzi N, Mancardi G, and Ferrarini M. Maintenance of B lymphocyte-related clones in the cerebrospinal fluid of multiple sclerosis patients. European journal of immunology. 2003;33(12):3433–8.

22. Corcione A, Casazza S, Ferretti E, Giunti D, Zappia E, Pistorio A, et al. Recapitulation of B cell differentiation in the central nervous system of patients with multiple sclerosis. Proceedings of the National Academy of Sciences of the United States of America. 2004;101(30):11064–9.

23. Cepok S, Rosche B, Grummel V, Vogel F, Zhou D, Sayn J, et al. SHort-Lived Plasma Blasts Are The Main B Cell Effector Subset During The Course Of Multiple Sclerosis. Brain : a journal of neurology. 2005;128(Pt 7):1667–76.

24. Taylor JJ, Pape KA, and Jenkins MK. A germinal center-independent pathway generates unswitched memory B cells early in the primary response. The Journal of experimental medicine. 2012;209(3):597–606.

25. Jones DD, Wilmore JR, and Allman D. Cellular Dynamics of Memory B Cell Populations: IgM+ and IgG+ Memory B Cells Persist Indefinitely as Quiescent Cells. Journal of immunology (Baltimore, Md: 1950). 2015; 195(10):4753–9.

26. Magliozzi R, Howell O, Vora A, Serafini B, Nicholas R, Puopolo M, et al. Meningeal B-cell follicles in secondary progressive multiple sclerosis associate with early onset of disease and severe cortical pathology. Brain : a journal of neurology. 2007;130(Pt 4):1089–104.

27. Confavreux C, Chapuis-Cellier C, Arnaud P, Robert O, Aimard G, and Devic M. Oligoclonal “fingerprint” of CSF IgG in multiple sclerosis patients is not modified following intrathecal administration of natural beta-interferon. Journal of neurology, neurosurgery, and psychiatry. 1986;49(11): 1308–12.

28. Li R, and Bar-Or A. The Multiple Roles of B Cells in Multiple Sclerosis and Their Implications in Multiple Sclerosis Therapies. Cold Spring Harbor perspectives in medicine. 2018.

29. Pitzalis C, Jones GW, Bombardieri M, and Jones SA. Ectopic lymphoid-like structures in infection, cancer and autoimmunity. Nature reviews Immunology. 2014;14(7):447–62.

30. Lovato L, Willis SN, Rodig SJ, Caron T, Almendinger SE, Howell OW, et al. Related B cell clones populate the meninges and parenchyma of patients with multiple sclerosis. Brain : a journal of neurology. 2011; 134(2):534–41.

31. Frischer JM, Bramow S, Dal-Bianco A, Lucchinetti CF, Rauschka H, Schmidbauer M, et al. The relation between inflammation and neurodegeneration in multiple sclerosis brains. Brain. 2009;132(Pt 5): 1175–89.

32. von Budingen HC, Bischof A, Eggers EL, Wang S, Bevan CJ, Cree BA, et al. Onset of secondary progressive MS after long-term rituximab therapy - a case report. Ann Clin Transl Neurol. 2017;4(1):46–52.

33. University of California SFMSET, Cree BA, Gourraud PA, Oksenberg JR, Bevan C, Crabtree-Hartman E, et al. Long-term evolution of multiple sclerosis disability in the treatment era. Annals of neurology. 2016;80(4):499–510.

34. Komori M, Lin YC, Cortese I, Blake A, Ohayon J, Cherup J, et al. Insufficient disease inhibition by intrathecal rituximab in progressive multiple sclerosis. Ann Clin Transl Neurol. 2016;3(3): 166–79.

35. Baker D, Marta M, Pryce G, Giovannoni G, and Schmierer K. Memory B Cells are Major Targets for Effective Immunotherapy in Relapsing Multiple Sclerosis. EBioMedicine. 2017;16:41–50.

36. Roll P, Palanichamy A, Kneitz C, Dorner T, and Tony HP. Regeneration of B cell subsets after transient B cell depletion using anti-CD20 antibodies in rheumatoid arthritis. Arthritis and rheumatism. 2006;54(8):2377–86.

37. Greenfield AL, and Hauser SL. B-cell Therapy for Multiple Sclerosis: Entering an era. Annals of neurology. 2018;83(1): 13–26.

38. Colucci M, Carsetti R, Cascioli S, Casiraghi F, Perna A, Ravà L, et al. B Cell Reconstitution after Rituximab Treatment in Idiopathic Nephrotic Syndrome. Journal of the American Society of Nephrology : JASN. 2016;27(6):1811–22.

39. Magliozzi R, Columba-Cabezas S, Serafini B, and Aloisi F. Intracerebral expression of CXCL13 and BAFF is accompanied by formation of lymphoid follicle-like structures in the meninges of mice with relapsing experimental autoimmune encephalomyelitis. J Neuroimmunol. 2004; 148(1-2): 11–23.

40. Serafini B, Rosicarelli B, Magliozzi R, Stigliano E, and Aloisi F. Detection of ectopic B-cell follicles with germinal centers in the meninges of patients with secondary progressive multiple sclerosis. Brain pathology (Zurich, Switzerland). 2004;14(2):164–74.

41. Lehmann-Horn K, Wang SZ, Sagan SA, Zamvil SS, and von Büdingen HC. B cell repertoire expansion occurs in meningeal ectopic lymphoid tissue. JCI Insight. 2016;1(20):e87234.

42. Horns F, Vollmers C, Croote D, Mackey SF, Swan GE, Dekker CL, et al. Lineage tracing of human B cells reveals the in vivo landscape of human antibody class switching. eLife. 2016;5:e16578.

43. Seifert M, Przekopowitz M, Taudien S, Lollies A, Ronge V, Drees B, et al. Functional capacities of human IgM memory B cells in early inflammatory responses and secondary germinal center reactions. Proceedings of the National Academy of Sciences. 2015;112(6):E546–E55.

44. Bar-Or A, Oliveira EM, Anderson DE, Krieger JI, Duddy M, O’Connor KC, et al. Immunological memory: contribution of memory B cells expressing costimulatory molecules in the resting state. J Immunol. 2001;167(10):5669–77.

45. Abramowski P, Otto B, and Martin R. The orally available, synthetic ether lipid edelfosine inhibits T cell proliferation and induces a type I interferon response. PLoS One. 2014;9(3):e91970.

46. Abbott NJ, Pizzo ME, Preston JE, Janigro D, and Thorne RG. The role of brain barriers in fluid movement in the CNS: is there a ‘glymphatic’ system? Acta Neuropathologica. 2018;135(3):387–407.

47. Engelhardt B, Carare RO, Bechmann I, Flügel A, Laman JD, and Weller RO. Vascular, glial, and lymphatic immune gateways of the central nervous system. Acta Neuropathologica. 2016;132:317–38.

48. Kappos L, Butzkueven H, Wiendl H, Spelman T, Pellegrini F, Chen Y, et al. Greater sensitivity to multiple sclerosis disability worsening and progression events using a roving versus a fixed reference value in a prospective cohort study. Multiple Sclerosis Journal. 2017;24(7):963–73.

49. McDonald WI, Compston A, Edan G, Goodkin D, Hartung HP, Lublin FD, et al. Recommended diagnostic criteria for multiple sclerosis: guidelines from the International Panel on the diagnosis of multiple sclerosis. Annals of neurology. 2001;50(1):121–7.

50. Bolotin DA, Poslavsky S, Mitrophanov I, Shugay M, Mamedov IZ, Putintseva EV, et al. MiXCR: software for comprehensive adaptive immunity profiling. Nat Methods. 2015; 12(5):380–1.

51. Hamming RW. Error Detecting and Error Correcting Codes. Bell System Technical Journal. 1950;29(2):147–60.

52. Krzywinski MI, Schein JE, Birol I, Connors J, Gascoyne R, Horsman D, et al. Circos: An information aesthetic for comparative genomics. Genome Research. 2009.

53. Giudicelli V, Brochet X, and Lefranc MP. IMGT/V-QUEST: IMGT standardized analysis of the immunoglobulin (IG) and T cell receptor (TR) nucleotide sequences. Cold Spring Harbor protocols. 2011;2011(6):695–715.

