## Supplementary material for "Longitudinally persistent cerebrospinal fluid B-cells resist treatment in multiple sclerosis"

**Supplementary Materials:**

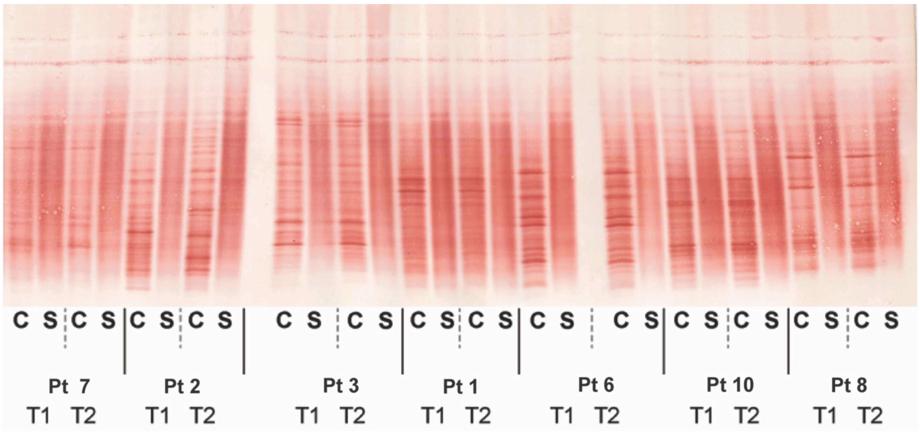


**Figure S1. CSF-unique OCBs are mostly stable over time.** Isoelectric focusing with IgG immunoblotting of CSF and serum at both time points for Pts 1-3, 6-8, and 10. The contrast and brightness of the original image were adjusted to improve visibility of bands, and the original image was flipped to present T1 and T2 in order and show CSF (C) before serum (S). The image was cropped to remove image parts that did not contain data relevant to this study. No specific features were enhanced, obscured, moved, removed, or introduced. The fact that patients do not appear in order is due to the order in which the respective samples were applied to the isoelectric focusing gel. Pt, patient. CSF (C), cerebrospinal fluid. (S), serum. OCB, oligoclonal band. Pt, patient. IgG, immunoglobulin G. T1, time point 1. T2, time point 2. Refer to table S2 for description of CSF OCB comparisons between T1 and T2.

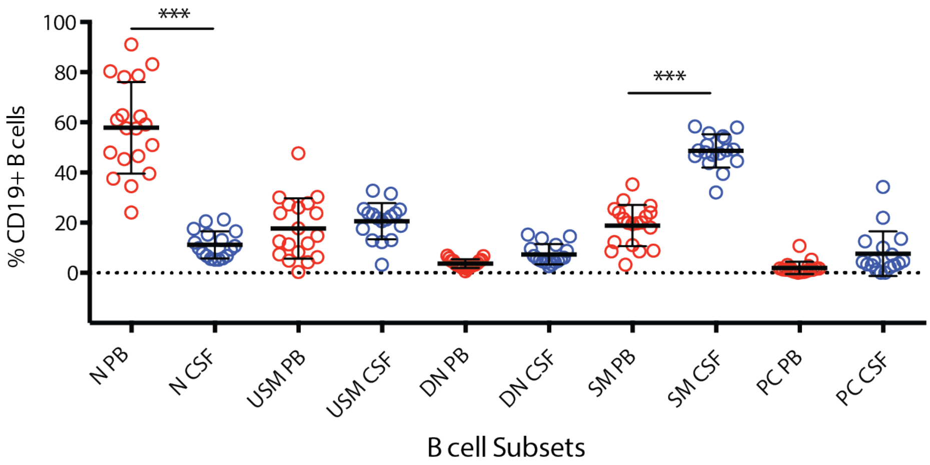


**Figure S2: Naïve B-cells are more prevalent in blood; in CSF, SM B-cells are relatively increased.** Shown are proportions of B-cell subsets in CSF (blue) and PB (red) among CD19+ B-cells as determined by multiparameter flow cytometry for N, USM, SM, DN, PC B-cell subsets. Overall CD19+ B-cells were 3.0% (+/- 3.0 SD) of all CSF lymphocytes and 7.2% (+/- 4.4 SD) of all PB lymphocytes. There were no significant differences between each subset per time point (not shown); therefore, shown here are combined data per subset from T1 and T2. T1-CSF subsets were measured in n=8 patients, in T2-CSF in n=9 patients, in T1-PB in n=9 patients, and in T2-PB in all 10 patients. Shown are naïve B-cells (N: CD19+IgD+CD27-), unswitched memory B-cells (USM: CD19+IgD+CD27+), class-switched memory B-cells (SM: CD19+IgD-CD27+), double negative B-cells (DN: CD19+IgD-CD27-), plasma cell (PC: CD27+CD38+ of CD19+IgD-), and CSF plasmablast/plasma cells (PC: CD19+IgD-CD27$^{hi}$). Comparisons between CSF and PB subsets were made using Anova (corrected for multiple comparisons using Sidak method); only significant differences are indicated, *** $p < 0.001$.

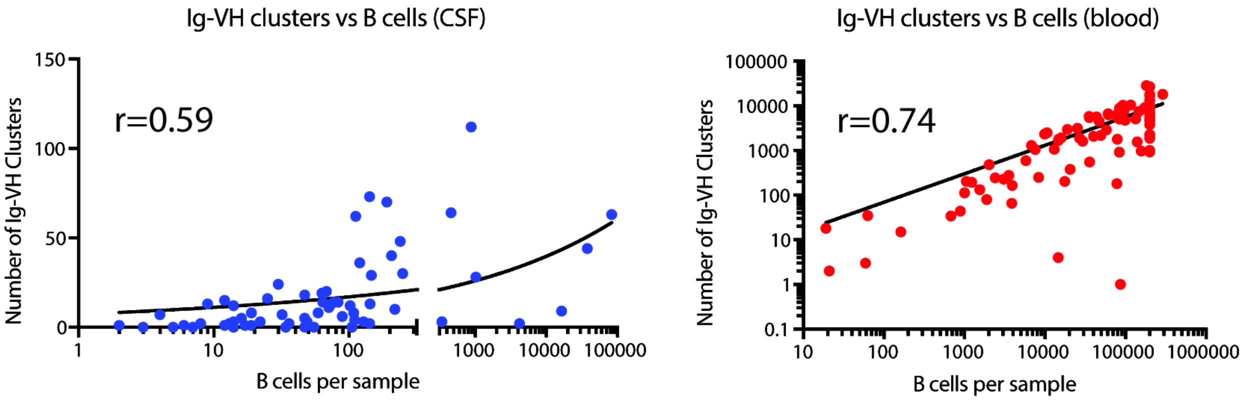


**Figure S3. Number of Ig-VH clusters in a sample is correlated with cell count.** Spearman correlation for number of Ig-VH clusters versus CSF B-cell count ($p<0.0001$) and for number of Ig-VH clusters versus PB B-cell count ($p<0.0001$) (log10 scale on x-axes as well as y-axis of PB plot). Ig-VH, immunoglobulin heavy chain variable region.

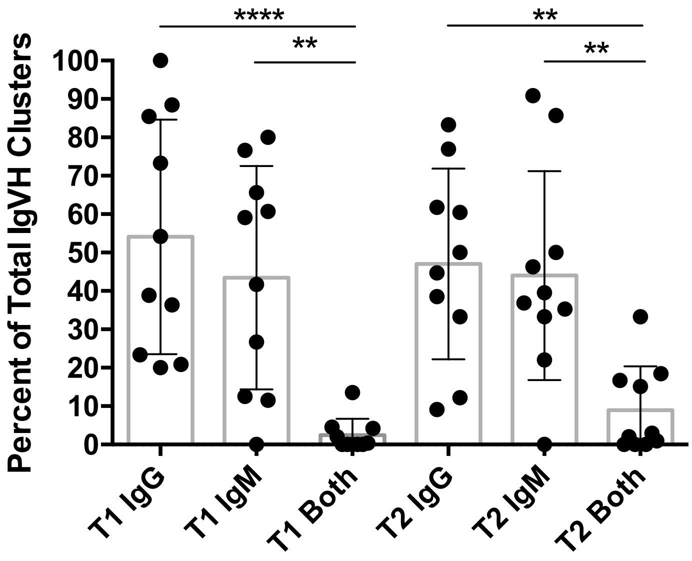


**Figure S4. The majority of CSF immune repertoire Ig-VH clusters express either IgM or IgG.** Each patient is represented by a point within each box plot showing the percentage of IgG-VH-only clusters or IgM-VH-only clusters, and of clusters with both IgG-VH and IgM-VH at T1 and T2. IgM, immunoglobulin M. Comparisons between Ig-VH cluster isotypes were made using Anova (corrected using Sidak method for multiple comparisons) in GraphPad Prism; only significant differences are indicated, ** $p < 0.01$, **** $p < 0.0001$.

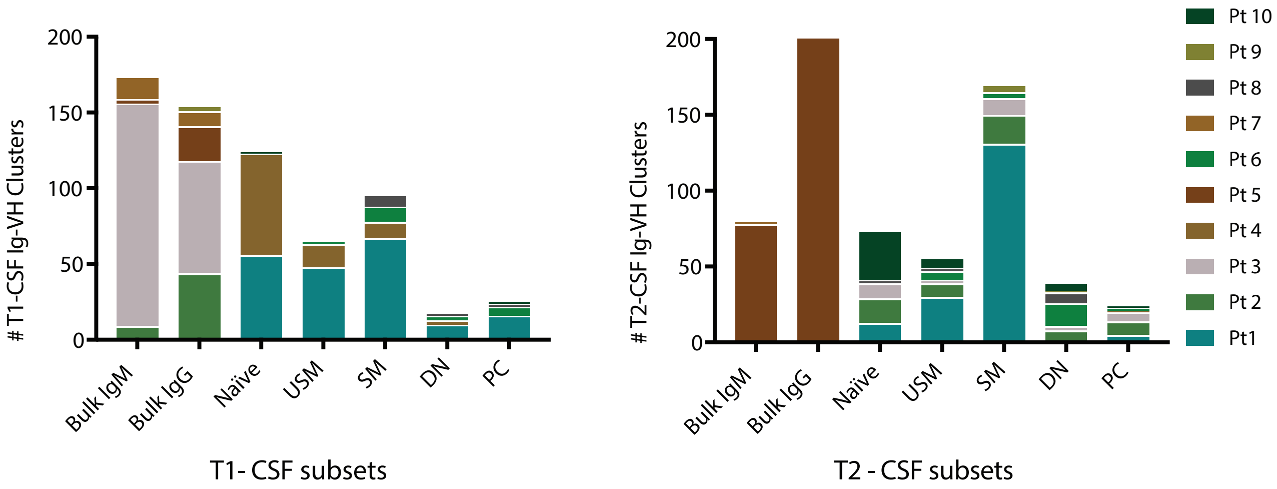


**Figure S5. Different B-cell subsets compose CSF Ig-VH repertoires at T1 and T2.** Of patients with sorted CSF B-cells, the number of Ig-VH clusters in T1-CSF and T2-CSF (and not in PB) containing each B-cell subset. As Ig-VH cluster is used as a unit of clonally-related populations in this study, this figure shows in which B-cell subsets these Ig-VH clusters have members. Bulk, unsorted B-cells.

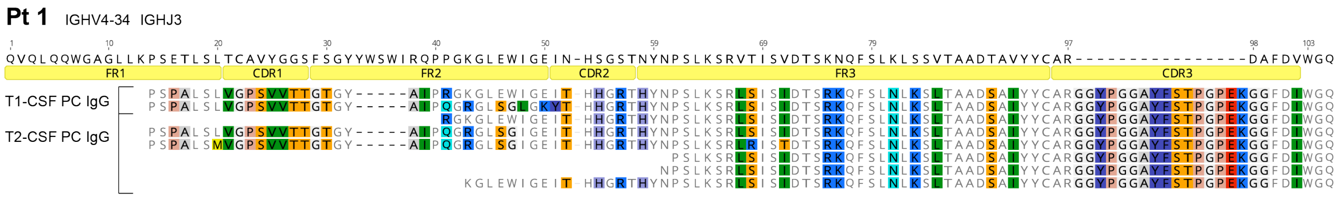


**Figure S6. Amino acid alignment of a CSF-persistent Ig-VH cluster.** Amino acid alignment of representative sequences from the indicated B-cell subsets from patient 1 together with the closest related germline IGHV. Color-shaded amino acids indicate differences from the germline. Regions of the immunoglobulin sequence are numbered and labeled according to IMGT (53) . Segments of the immunoglobulin sequence are labeled: FR1-FR3, framework regions 1-3; CDR1-CDR3, complementarity determining regions 1-3. Alignments generated using Geneious and IMGT High-V Quest. IGHV, immunoglobulin heavy chain variable germline segment. IGHJ, immunoglobulin heavy chain joining germline segment.

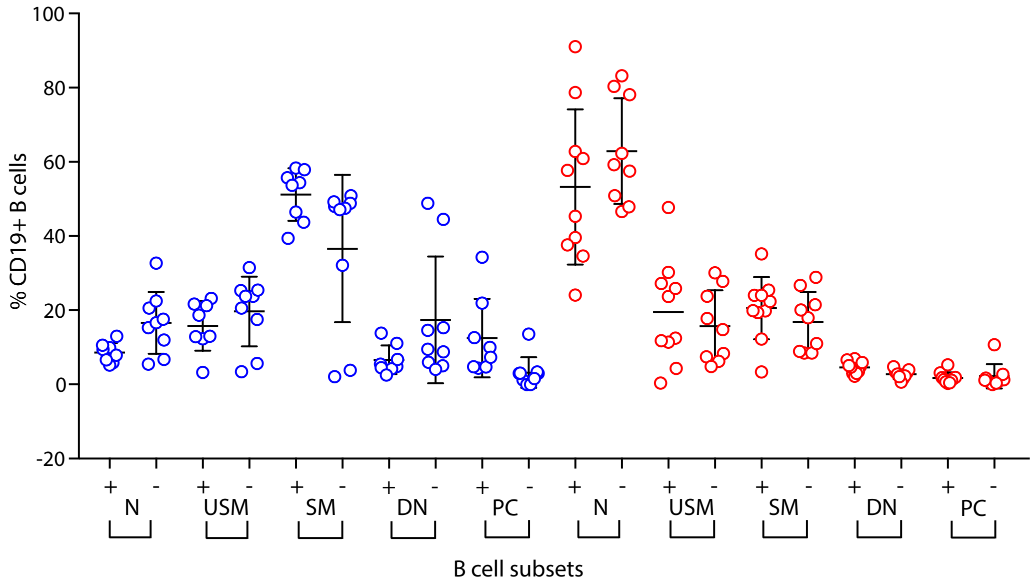


**Figure S7. Patients with persistent CSF Ig-VH clusters show no significant difference in B-cell type prevalence in CSF or PB compared to patients without persistent CSF Ig-VH clusters.** Blue circles: CSF. Red circles: PB. Shown are naïve B-cells (N: CD19+IgD+CD27-), unswitched memory B-cells (USM: CD19+IgD+CD27+), class-switched memory B-cells (SM: CD19+IgD-CD27+), double negative B-cells (DN: CD19+IgD-CD27-), plasma cell (PC: CD27+CD38+ of CD19+IgD-), and CSF plasmablast/plasma cells (PC: CD19+IgD-CD27$^{hi}$). (+), patients with persistent CSF Ig-VH clusters. (-), patients without persistent CSF Ig-VH clusters. T1-CSF subsets were measured in n=8 patients, in T2-CSF in n=9 patients, in T1-PB in n=9 patients, and in T2-PB in all 10 patients. Kruskal-Wallis with Dunn correction for multiple comparisons, $p<0.05$ was considered significant (none of the (+) vs (-) comparisons were statistically significant).

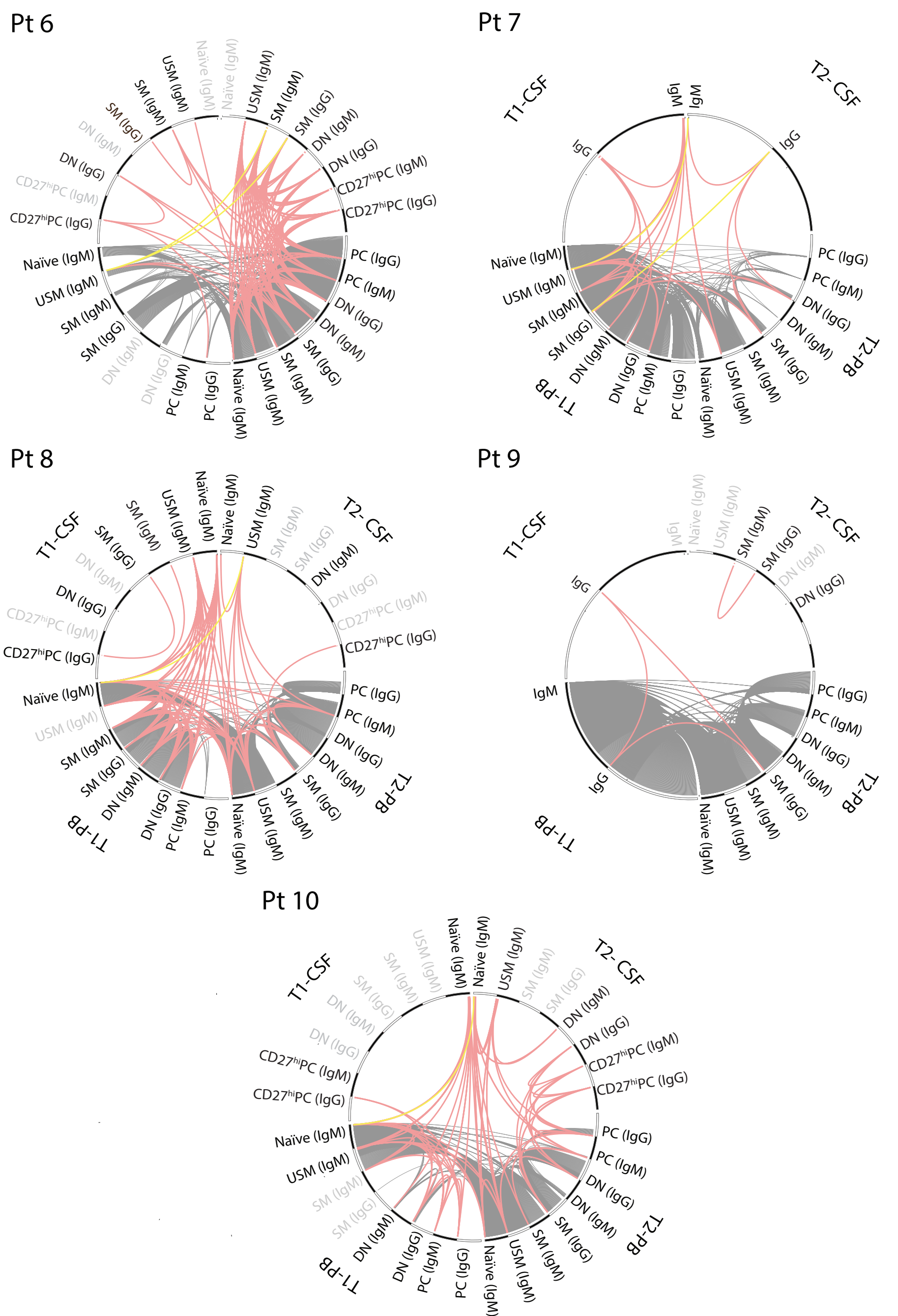


**Figure S8. Patients without identifiable persistent CSF Ig-VH clusters still have clonal connections between CSF and PB.** Within T1, Ig-VH clusters spanning CSF and PB are often SM, followed closely by USM. At T2, CSF-PB Ig-VH clusters are often SM. Clonal relationships between B-cell subsets are shown for each patient. Lines represent Ig-VH clusters shared between two subsets. Grey lines: PB-only Ig-VH clusters. Red lines: CSF-containing Ig-VH clusters. Yellow lines: T1-PB B-cell subsets, or bulk PB IgG and IgM (from Pt 9), that provide input to T2-CSF without involving T1-CSF. Grey font indicates subsets/Ig isotypes from which no Ig-VH libraries could be obtained.

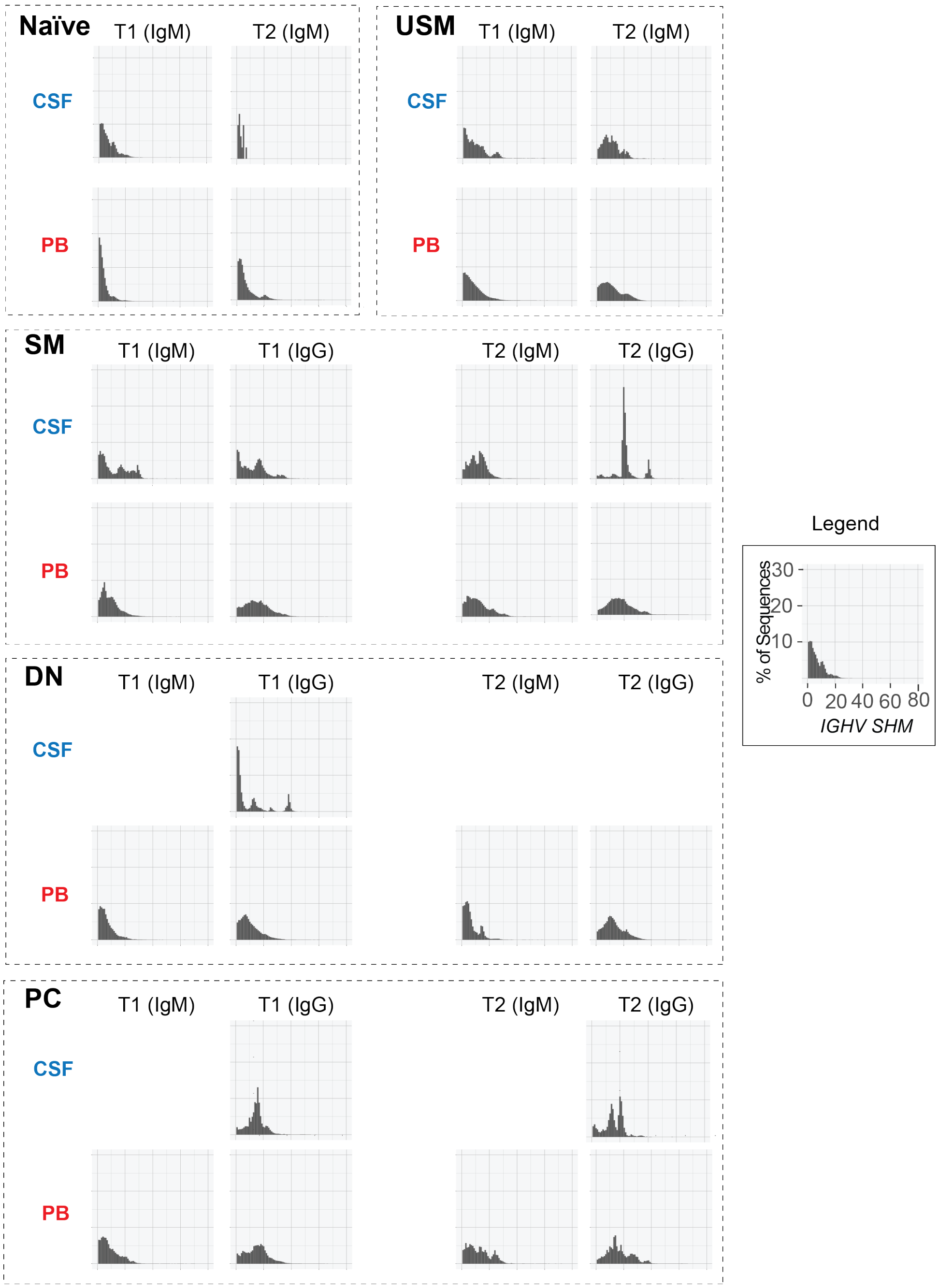


**Figure S9. Somatic hypermutation rates follow expected patterns along B-cell lineage.** Shown are somatic hypermutation profiles for B-cell subsets in CSF and PB from Patient 1. The x-axis shows the number of amino acid differences from reference germline IGHV sequences, i.e. mutations. The y-axis shows the percentage of sequences in the sample with a given number of mutations on the x-axis. Overall, the degree of somatic hypermutation follows the expected increase along the B-cell maturation stages as antigen exposure and affinity maturation occur: somatic hypermutation is least in naïve B-cells, and greater in IgG-expressing SM and PC. In this patient, there is a particularly high degree of SHM in IgG SM B-cells in T2-CSF. SHM, somatic hypermutation.

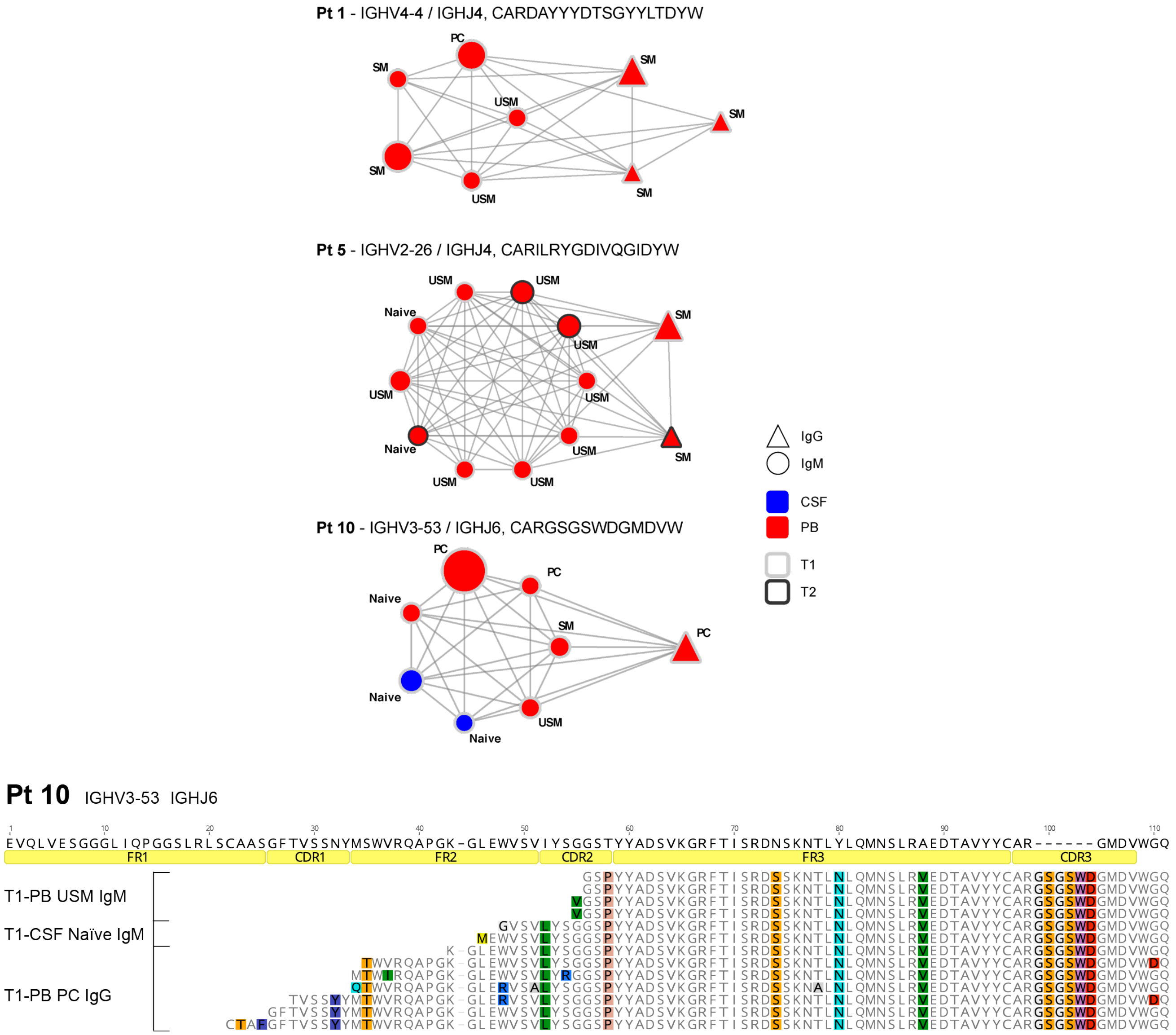


**Figure S10: Clonal relationships between IgM-expressing USM B-cells and IgG-expressing B-cell subsets suggest Ig class-switch recombination and further maturation of USM B-cells.** Shown are three representative Ig-VH cluster networks of clonally related B-cells, with IGHV, IGHJ and most common CDR3 amino acid sequences indicated per network. Each node represents a specific CDR3 expressed by the indicated B-cell subset; the node size is relative to the number of sequences found for each IGHV-IGHJ-CDR3 combination (range 2 to 317,112). IgM-expressing B-cell subsets are represented by circles, those expressing IgG by triangles. CSF B-cell subsets are indicated in blue, PB subsets in red; light gray rims indicate T1 subsets, dark gray rims indicated subsets that derive from T2. Shown below the cluster networks is an amino acid alignment of representative sequences from the indicated B-cell subsets from patient 10 together with the closest related germline IGHV. Color-shaded amino acids indicate differences from the germline. Regions of the immunoglobulin sequence are numbered and labeled according to IMGT (53).

**Table S1. Patient characteristics based on presence or absence of CSF persistent Ig-VH**

| | Patients with persistent CSF Ig-VH clusters (n=5) | Patients without persistent CSF Ig-VH clusters (n=5) |
|---|---|---|
| **Age (years)** | 30.4 (+/-3.1) | 40 (+/-3.8) |
| **Sex (M, F)*** | 0, 5 | 4, 1 |
| **Disease Duration (years)** | 2.2 (+/-1.2) | 1.4 (+/-0.6) |
| **EDSS** | 3.4 (+/-0.5) | 2.7 (+/-0.8) |
| **Time between T1 and T2 (months)** | 15.2 (+/- 1.8) | 13 (+/- 1.1) |
| **Clinical relapse between T1-T2** | 4 | 2 |
| **IgG Index (normal <0.66)** | 1.2 (+/-0.1) | 0.9 (+/-0.1) |
| **No. of Patients on IMT** | 4 | 3 |
| **Anti-lymphocyte trafficking IMT (i.e. fingolimod or natalizumab)** | 2 | 2 |
| **Gadolinium enhancement on MRI (at T1, T2)** | 4, 2 | 4, 2 |
| **CSF volume (mL)** | 11.5 (+/- 3.9) | 13.7 (+/- 3.3) |

Patients with and without persistent CSF Ig-VH clusters did not differ with respect to clinical metrics. More men had persistent CSF B-cells than women (*$p<0.05$ Fisher's exact test). EDSS, expanded disability status scale. IMT, immune modulating therapy. MRI, magnetic resonance imaging. IgG index, immunoglobulin G index ((CSF IgG/CSF albumin)/(serum IgG/serum albumin)).

**Table S2. Clinical CSF biometrics.**

| Patient ID | Time Point | CSF WBC (cells/uL) | CSF collected (mL) | IgG Index (normal <0.66) | OCB | OCB Comparison |
|---|---|---|---|---|---|---|
| **1** | T1<br>T2 | 8<br>3 | 15<br>16 | 0.83<br>0.65 | 5<br>3 | Decrease in number |
| **2** | T1<br>T2 | 12<br>10 | 6<br>10 | 1.1<br>1.75 | >5<br>>5 | Increase in number |
| **3** | T1<br>T2 | 7<br>3 | 7<br>17.5 | NP<br>0.75 | >5<br>>5 | Stable |
| **4** | T1<br>T2 | 3<br>1 | 9.5<br>10 | 1.27<br>0.94 | >5<br>>5 | NP |
| **5** | T1<br>T2 | 4<br>4 | 14<br>10 | 1.64<br>1.55 | >5<br>>5 | NP |
| **6** | T1<br>T2 | 3<br>1 | 16<br>17 | 1<br>0.85 | 5<br>>5 | Stable |
| **7** | T1<br>T2 | 2<br>1 | 9.5<br>11.5 | 0.9<br>0.74 | >5<br>5 | Stable |
| **8** | T1<br>T2 | 4<br>2 | 14<br>17 | 0.62<br>0.67 | >5<br>>5 | Stable overall: 1 band more prominent, 1 less prominent |
| **9** | T1<br>T2 | 0<br>0 | 9.5<br>13.5 | 1<br>0.62 | >5<br>>5 | NP |
| **10** | T1<br>T2 | 8<br>12 | 11<br>10 | 1.48<br>1.32 | >5<br>>5 | Stable overall: 1 band more prominent |

Clinical diagnostic laboratory CSF WBC count, IgG index and OCBs present are shown for each patient at each time point. In n=7 patients, there was additional available CSF, and in these patients the pattern of CSF OCBs at T2 was compared to the OCB pattern at T1. NP, not performed. WBC, white blood cell. IgG index, immunoglobulin G index ((CSF IgG/CSF albumin)/(serum IgG/serum albumin)). OCB, oligoclonal band.

### Table S3. B-cell samples analyzed by IgSeq.

| Pt ID | Time Point | Sample Type | B-cell Subset | Number of B-cells | Isotype | Ig-VH Clusters | Exp | Raw Reads | Aligned Reads | |
|---|---|---|---|---|---|---|---|---|---|---|
| 1 | T1 | CSF | Naïve | 112 | IgM | 62 | I | 39948 | 20587 | |
| | | | USM | 142 | IgM | 73 | I | 49025 | 24978 | |
| | | | SM | 460 | IgG | 64 | I | 53241 | 23042 | |
| | | | | | IgM | 35 | I | 9615 | 5255 | |
| | | | DN | 83 | IgG | 14 | I | 37097 | 12231 | |
| | | | | | IgM | 0 | I | 0 | 0 | |
| | | | PC | 120 | IgG | 36 | I | 207795 | 86877 | |
| | | | | | IgM | 0 | I | 0 | 0 | |
| | | PB | Naïve | 175179 | IgM | 9151 | I | 220760 | 41385 | |
| | | | USM | 84463 | IgM | 8832 | I | 339153 | 105179 | |
| | | | SM | 95720 | IgG | 7842 | I | 553275 | 180184 | |
| | | | | | IgM | 635 | I | 219349 | 103118 | |
| | | | DN | 19072 | IgG | 2963 | I | 626941 | 234604 | |
| | | | | | IgM | 99 | I | 73403 | 12421 | |
| | | | PC | 1055 | IgG | 202 | I | 846488 | 298704 | |
| | | | | | IgM | 124 | I | 195769 | 81500 | |
| | T2 | CSF | Naïve | 77 | IgM | 14 | I | 659 | 0 | |
| | | | | | | | TR | 9456 | 1420 | |
| | | | | | | | TR | 162 | 0 | |
| | | | USM | 239 | IgM | 48 | I | 157 | 0 | |
| | | | | | | | TR | 26 | 0 | |
| | | | | | | | TR | 2989 | 1009 | |
| | | | SM | 880 | IgG | 112 | I | 142 | 67 | |
| | | | | | | | TR | 161 | 0 | |
| | | | | | | | TR | 5871 | 2323 | |
| | | | | | | | TR | 1332980 | 625014 | |
| | | | | | IgM | 83 | I | 59 | 0 | |
| | | | | | | | TR | 285201 | 159982 | |
| | | | | | | | TR | 192 | 13 | |
| | | | | | | | TR | 0 | 0 | |
| | | | DN | 47 | IgG | 0 | I | 339 | 0 | |
| | | | | | | | TR | 1325 | 0 | * |
| | | | | | IgM | 0 | I | 6480 | 0 | |
| | | | | | | | TR | 2143 | 0 | |
| | | | PC | 71 | IgG | 11 | I | 736 | 201 | |
| | | | | | | | TR | 330 | 0 | |
| | | | | | | | TR | 11873 | 4531 | |
| | | | | | IgM | 0 | I | 3 | 0 | |
| | | | | | | | TR | 0 | 0 | |
| | | | | | | | TR | 77 | 0 | |
| | | PB | Naïve | 60917 | IgM | 6640 | I | 262400 | 49100 | |
| | | | USM | 79605 | IgM | 5480 | I | 266747 | 53849 | |
| | | | SM | 47200 | IgG | 4482 | I | 769897 | 220884 | |
| | | | | | IgM | 1101 | I | 231100 | 56482 | |

| | | | | | | | | | |
|---|---|---|---|---|---|---|---|---|---|
| | | | DN | 14654 | IgG | 1675 | I | 609380 | 185738 |
| | | | | | IgM | 97 | I | 46488 | 12816 |
| | | | PC | 1229 | IgG | 194 | I | 550130 | 190036 |
| | | | | | IgM | 125 | I | 139235 | 50504 |
| **2** | T1 | CSF | bulk | 37673 | IgG | 44 | I | 81628 | 45243 |
| | | | | | IgM | 7 | I | 1133 | 308 |
| | | PB | Naïve | 200000 | IgM | 7822 | I | 152323 | 31911 |
| | | | USM | 146293 | IgM | 7450 | I | 162622 | 47212 |
| | | | SM | 200000 | IgG | 6793 | I | 423906 | 155698 |
| | | | | | IgM | 1117 | I | 81672 | 37788 |
| | | | DN | 43729 | IgG | 5654 | I | 577341 | 244324 |
| | | | | | IgM | 650 | I | 172128 | 95214 |
| | | | PC | 7585 | IgG | 1049 | I | 349470 | 149936 |
| | | | | | IgM | 211 | I | 50291 | 28296 |
| | T2 | CSF | Naïve | 47 | IgM | 18 | I | 33126 | 17856 |
| | | | USM | 102 | IgM | 12 | I | 32198 | 19097 |
| | | | SM | 249 | IgG | 30 | I | 92267 | 32018 |
| | | | | | IgM | 11 | I | 2486 | 569 |
| | | | DN | 30 | IgG | 24 | I | 46805 | 19529 |
| | | | | | IgM | 0 | I | 34 | 0 |
| | | | PC | 68 | IgG | 20 | I | 50993 | 23903 |
| | | | | | IgM | 0 | I | 2151 | 1 |
| | | PB | Naïve | 61533 | IgM | 6308 | I | 154660 | 39006 |
| | | | USM | 10634 | IgM | 2508 | I | 146168 | 57403 |
| | | | SM | 15881 | IgG | 1926 | I | 562184 | 184976 |
| | | | | | IgM | 285 | I | 83634 | 30364 |
| | | | DN | 6787 | IgG | 1297 | I | 337274 | 150492 |
| | | | | | IgM | 85 | I | 29104 | 15681 |
| | | | PC | 63 | IgG | 35 | I | 359362 | 114119 |
| | | | | | IgM | 8 | I | 107154 | 66322 |
| **3** | T1 | CSF | bulk | 82900 | IgG | 63 | I | 1161091 | 522507 |
| | | | | | IgM | 140 | I | 209102 | 39341 |
| | | PB | Naïve | 200000 | IgM | 9660 | I | 745241 | 161869 |
| | | | USM | 84400 | IgM | 5030 | I | 732578 | 209200 |
| | | | SM | 200000 | IgG | 925 | I | 1163560 | 373157 |
| | | | | | IgM | 365 | I | 352938 | 111239 |
| | | | DN | 170000 | IgG | 8572 | I | 538099 | 150224 |
| | | | | | IgM | 783 | I | 109361 | 25553 |
| | | | PC | 2410 | IgG | 245 | I | 985465 | 379640 |
| | | | | | IgM | 179 | I | 223450 | 100665 |
| | T2 | CSF | Naïve | 14 | IgM | 12 | I | 635998 | 196072 |
| | | | USM † | 4 | IgM | 7 | I | 2083 | 172 |
| | | | SM | 63 | IgG | 19 | I | 875008 | 368535 |

| | | | | | | | | | | |
|---|---|---|---|---|---|---|---|---|---|---|
| | | | | | IgM | 0 | I | 291842 | 0 | |
| | | | DN | 17 | IgG | 1 | I | 1036 | 4 | |
| | | | | | IgM | 4 | I | 2311 | 114 | |
| | | | PC † | 9 | IgG | 13 | I | 666444 | 402357 | |
| | | | | | | | TR | 600838 | 492251 | |
| | | | | | IgM | 1 | I | 27352 | 1 | |
| | | | | | | | TR | 35400 | 3 | |
| | | PB | Naïve | 200000 | IgM | 10223 | I | 461166 | 66739 | |
| | | | USM | 14878 | IgM | 1818 | I | 394218 | 124183 | |
| | | | SM | 133478 | IgG | 5088 | I | 830086 | 315358 | |
| | | | | | IgM | 459 | I | 114407 | 34347 | |
| | | | DN | 92953 | IgG | 10286 | I | 1548667 | 670757 | |
| | | | | | IgM | 854 | I | 287615 | 88721 | |
| | | | PC | 13038 | IgG | 1062 | I | 737288 | 330353 | |
| | | | | | IgM | 595 | I | 208666 | 62988 | |
| **4** | T1 | CSF | Naïve | 190 | IgM | 70 | I | 33043 | 13483 | |
| | | | USM | 146 | IgM | 29 | I | 24853 | 8364 | |
| | | | SM-PC | 1022 | IgG | 28 | I | 53400 | 18138 | |
| | | | | | IgM | 0 | I | 0 | 0 | |
| | | | DN | 89 | IgG | 6 | I | 86651 | 12354 | |
| | | | | | IgM | 0 | I | 0 | 0 | |
| | | PB | Naïve | 290198 | IgM | 18074 | I | 466054 | 97254 | |
| | | | USM | 89486 | IgM | 7123 | I | 415706 | 125276 | |
| | | | SM | 188264 | IgG | 8258 | I | 1160979 | 249186 | |
| | | | | | IgM | 1081 | I | 245366 | 78639 | |
| | | | DN | 35706 | IgG | 5492 | I | 1912127 | 469120 | |
| | | | | | IgM | 253 | I | 279055 | 60085 | |
| | | | PC | 1001 | IgG | 113 | I | 1008112 | 207739 | |
| | | | | | IgM | 50 | I | 89606 | 41889 | |
| | T2 | CSF | Naïve | 19 | IgM | 1 | I | 16961 | 17 | |
| | | | | | | | TR | 2764 | 0 | |
| | | | USM | 34 | IgM | 0 | I | 83755 | 0 | * |
| | | | | | | | TR | 50290 | 0 | |
| | | | SM | 142 | IgG | 2 | I | 1863 | 1045 | |
| | | | | | | | TR | 18520 | 8543 | |
| | | | | | | | TR | 723 | 51 | |
| | | | | | IgM | 0 | I | 3 | 0 | |
| | | | | | | | TR | 2 | 0 | |
| | | | | | | | TR | 31 | 0 | |
| | | | DN | 14 | IgG | 0 | I | 622 | 0 | |
| | | | | | | | TR | 106 | 0 | * |
| | | | | | IgM | 0 | I | 4831 | 0 | |
| | | | | | | | TR | 764 | 0 | |

| | | | | | | | | | |
|---|---|---|---|---|---|---|---|---|---|
| | | | PC | 12 | IgG | 1 | I | 2266 | 559 |
| | | | | | | | TR | 19903 | 10698 |
| | | | | | | | TR | 675 | 0 |
| | | | | | IgM | 1 | I | 510 | 0 |
| | | | | | | | TR | 16334 | 11 |
| | | | | | | | TR | 189 | 0 |
| | | PB | Naïve | 200000 | IgM | 2107 | I | 163845 | 8893 |
| | | | USM | 200000 | IgM | 2133 | I | 140853 | 17597 |
| | | | SM | 200000 | IgG | 9447 | I | 514188 | 138860 |
| | | | | | IgM | 541 | I | 53707 | 6956 |
| | | | DN | 40109 | IgG | 2082 | I | 584643 | 166918 |
| | | | | | IgM | 474 | I | 81251 | 14395 |
| | | | PC | 679 | IgG | 34 | I | 282182 | 92666 |
| | | | | | IgM | 59 | I | 203153 | 51466 |
| **5** | T1 | CSF | bulk | unknown | IgG | 23 | I | 1361 | 325 |
| | | | | | | | TR | 23091 | 7962 |
| | | | | | | | TR | 571 | 0 |
| | | | | | IgM | 3 | I | 132 | 0 |
| | | | | | | | TR | 2595 | 386 |
| | | | | | | | TR | 43 | 0 |
| | | PB | Naïve | 200000 | IgM | 4976 | I | 242268 | 49347 |
| | | | USM | 200000 | IgM | 4523 | I | 257151 | 59188 |
| | | | SM | 200000 | IgG | 1876 | I | 720826 | 249298 |
| | | | | | IgM | 253 | I | 123144 | 46742 |
| | | | DN | 81400 | IgG | 6827 | I | 746881 | 229740 |
| | | | | | IgM | 246 | I | 35456 | 7616 |
| | | | PC | 1890 | IgG | 80 | I | 223187 | 94706 |
| | | | | | IgM | 147 | I | 595747 | 270258 |
| | T2 | CSF | bulk | unknown | IgG | 245 | I | 14221 | 2631 |
| | | | | | | | TR | 1324791 | 564943 |
| | | | | | IgM | 70 | I | 260 | 3 |
| | | | | | | | TR | 14057 | 2843 |
| | | PB | Naïve | 200000 | IgM | 16802 | I | 366778 | 116156 |
| | | | USM | 200000 | IgM | 4907 | I | 414710 | 202128 |
| | | | SM | 200000 | IgG | 9098 | I | 896283 | 295044 |
| | | | | | IgM | 472 | I | 50416 | 15033 |
| | | | DN | 115849 | IgG | 10397 | I | 809315 | 258011 |
| | | | | | IgM | 148 | I | 19478 | 3128 |
| | | | PC | 162 | IgG | 15 | I | 543166 | 217384 |
| | | | | | IgM | 20 | I | 178884 | 77425 |
| **6** | T1 | CSF | Naïve | 105 | IgM | 0 | I | 264476 | 0 * |
| | | | USM | 128 | IgM | 3 | I | 327565 | 55744 |
| | | | SM | 340 | IgG | 3 | I | 31953 | 14499 |

| | | | | | | | | | |
|---|---|---|---|---|---|---|---|---|---|
| | | | | | IgM | 9 | I | 336665 | 158245 |
| | | | DN | 110 | IgG | 4 | I | 325684 | 10319 |
| | | | | | IgM | 0 | I | 32410 | 0 |
| | | | PC | 19 | IgG | 8 | I | 764076 | 122329 |
| | | | | | IgM | 0 | I | 8833 | 0 |
| | | PB | Naïve | 200000 | IgM | 1002 | I | 543933 | 5352 |
| | | | USM | 78176 | IgM | 180 | I | 257389 | 36782 |
| | | | SM | 156710 | IgG | 972 | I | 237251 | 13302 |
| | | | | | IgM | 199 | I | 170953 | 14970 |
| | | | DN | 35762 | IgG | 550 | I | 145676 | 3856 * |
| | | | | | IgM | 16 | I | 7339 | 167 |
| | | | PC | 889 | IgG | 44 | I | 394103 | 61993 |
| | | | | | IgM | 28 | I | 185208 | 28922 |
| | T2 | CSF | Naïve | 45 | IgM | 0 | I | 0 | 0 * |
| | | | USM | 64 | IgM | 14 | I | 396630 | 356366 |
| | | | SM | 143 | IgG | 13 | I | 506718 | 391468 |
| | | | | | IgM | 10 | I | 333972 | 260293 |
| | | | DN | 25 | IgG | 16 | I | 688318 | 521557 |
| | | | | | IgM | 5 | I | 112268 | 667 |
| | | | PC† | 12 | IgG | 15 | I | 470279 | 378332 |
| | | | | | IgM | 12 | I | 449574 | 349612 |
| | | PB | Naïve | 200000 | IgM | 17858 | I | 375931 | 182899 |
| | | | USM | 78954 | IgM | 1792 | I | 453932 | 325066 |
| | | | SM | 200000 | IgG | 5140 | I | 528853 | 365527 |
| | | | | | IgM | 891 | I | 151177 | 99882 |
| | | | DN | 98481 | IgG | 9108 | I | 668534 | 444711 |
| | | | | | IgM | 412 | I | 116891 | 77454 |
| | | | PC | 26941 | IgG | 1899 | I | 704335 | 489843 |
| | | | | | IgM | 860 | I | 237060 | 171077 |
| **7** | T1 | CSF | bulk | 16399 | IgG | 9 | I | 50692 | 15008 |
| | | | | | IgM | 15 | I | 762 | 324 |
| | | PB | Naïve | 200000 | IgM | 6026 | I | 152040 | 25046 |
| | | | USM | 200000 | IgM | 4731 | I | 159363 | 39758 |
| | | | SM | 200000 | IgG | 6839 | I | 437850 | 126749 |
| | | | | | IgM | 2096 | I | 132495 | 48575 |
| | | | DN | 86617 | IgG | 8579 | I | 429833 | 125093 |
| | | | | | IgM | 1440 | I | 80089 | 23517 |
| | | | PC | 5785 | IgG | 589 | I | 462064 | 166191 |
| | | | | | IgM | 405 | I | 93627 | 44582 |
| | T2 | CSF | bulk | 4210 | IgG | 2 | I | 35623 | 18926 |
| | | | | | IgM | 2 | I | 177 | 97 |
| | | PB | Naïve | 25417 | IgM | 3128 | I | 115211 | 45698 |
| | | | USM | 9912 | IgM | 2335 | I | 193174 | 84241 |

| | | | | | | | | | | |
|---|---|---|---|---|---|---|---|---|---|---|
| | | | SM | 3078 | IgG | 228 | I | 946058 | 364072 | |
| | | | | | IgM | 54 | I | 190029 | 76307 | |
| | | | DN | 1551 | IgG | 132 | I | 421432 | 157055 | |
| | | | | | IgM | 4 | I | 13520 | 1396 | |
| | | | PC † | 19 | IgG | 18 | I | 143120 | 49583 | |
| | | | | | IgM | 10 | I | 531331 | 220280 | |
| **8** | T1 | CSF | Naïve | 32 | IgM | 7 | I | 117980 | 76362 | |
| | | | USM | 59 | IgM | 8 | I | 121560 | 101569 | |
| | | | SM | 108 | IgG | 8 | I | 1627 | 867 | |
| | | | | | | | TR | 14353 | 5565 | |
| | | | | | | | TR | 380 | 0 | |
| | | | | | IgM | 1 | I | 31 | 0 | |
| | | | | | | | TR | 325 | 170 | |
| | | | | | | | TR | 6 | 0 | |
| | | | DN | 36 | IgG | 2 | I | 1412 | 234 | |
| | | | | | | | TR | 12793 | 4419 | |
| | | | | | | | TR | 375 | 0 | |
| | | | | 36 | IgM | 0 | I | 84 | 0 | |
| | | | | | | | TR | 1378 | 0 | |
| | | | | | | | TR | 23 | 0 | |
| | | | PC | 8 | IgG | 2 | I | 1524 | 0 | |
| | | | | | | | TR | 290934 | 300 | |
| | | | | | | | TR | 14464 | 45 | |
| | | | | | | | TR | 430 | 0 | |
| | | | | 8 | IgM | 0 | I | 1 | 0 | |
| | | | | | | | TR | 20 | 0 | |
| | | | | | | | TR | 334 | 0 | |
| | | | | | | | TR | 0 | 0 | |
| | | PB | Naïve | 181753 | IgM | 28341 | I | 3938585 | 1131870 | |
| | | | USM | 14649 | IgM | 4 | I | 107 | 30 | * |
| | | | SM | 17624 | IgG | 203 | I | 969745 | 423212 | |
| | | | | | IgM | 131 | I | 249957 | 96678 | |
| | | | DN | 3909 | IgG | 165 | I | 758428 | 267570 | |
| | | | | | IgM | 75 | I | 145441 | 46026 | |
| | | | PC | 21 | IgG | 2 | I | 168262 | 105243 | |
| | | | | | IgM | 5 | I | 37121 | 27987 | |
| | T2 | CSF | Naïve | 13 | IgM | 2 | I | 4724 | 1164 | |
| | | | USM | 22 | IgM | 3 | I | 8453 | 2550 | |
| | | | SM | 55 | IgG | 0 | I | 16 | 0 | |
| | | | | | IgM | 0 | I | 75 | 0 | * |
| | | | DN † | 7 | IgG | 0 | I | 4 | 0 | |
| | | | | | IgM | 15 | I | 9655 | 1999 | |
| | | | PC | 2 | IgG | 1 | I | 78520 | 32697 | |

| | | | | | | | | | | |
|---|---|---|---|---|---|---|---|---|---|---|
| | | | | | IgM | 0 | I | 27699 | 0 | |
| | | PB | Naïve | 200000 | IgM | 13265 | I | 551204 | 146660 | |
| | | | USM | 57800 | IgM | 2908 | I | 2020938 | 634198 | |
| | | | SM | 140118 | IgG | 1553 | I | 593767 | 182970 | |
| | | | | | IgM | 0 | I | 26 | 2 | |
| | | | DN | 20474 | IgG | 378 | I | 544174 | 160618 | |
| | | | | | IgM | 151 | I | 261438 | 87403 | |
| | | | PC | 3565 | IgG | 277 | I | 122810 | 38172 | |
| | | | | | IgM | 210 | I | 155272 | 55287 | |
| **9** | T1 | CSF | bulk | unknown | IgG | 4 | I | 1960 | 874 | |
| | | | | | | | TR | 700 | 0 | |
| | | | | | | | TR | 18527 | 6855 | |
| | | | | unknown | IgM | 0 | I | 59 | 0 | |
| | | | | | | | TR | 21 | 0 | |
| | | | | | | | TR | 1097 | 0 | |
| | | PB | bulk | unknown | IgG | 960 | I | 1017427 | 703518 | |
| | | | | | IgM | 10559 | I | 3698830 | 2163377 | |
| | T2 | CSF | Naïve | 3 | IgM | 0 | I | 25 | 0 | * |
| | | | USM | 5 | IgM | 0 | I | 20 | 0 | * |
| | | | SM | 16 | IgG | 5 | I | 17095 | 8150 | |
| | | | | | IgM | 1 | I | 34416 | 99 | |
| | | | DN | 6 | IgG | 1 | I | 9533 | 17 | |
| | | | | | IgM | 0 | I | 1031 | 0 | |
| | | | PC | 0 | IgG | n/a | n/a | n/a | n/a | |
| | | | | | IgM | n/a | n/a | n/a | n/a | |
| | | PB | Naïve | 200000 | IgM | 26685 | I | 463680 | 172023 | |
| | | | USM | 35263 | IgM | 5757 | I | 440115 | 253640 | |
| | | | SM | 98748 | IgG | 4733 | I | 818738 | 452511 | |
| | | | | | IgM | 289 | I | 79508 | 39915 | |
| | | | DN | 29358 | IgG | 1621 | I | 767356 | 487204 | |
| | | | | | IgM | 186 | I | 73377 | 21868 | |
| | | | PC | 2031 | IgG | 484 | I | 494349 | 309270 | |
| | | | | | IgM | 106 | I | 42157 | 26743 | |
| **10** | T1 | CSF | Naïve | 74 | IgM | 13 | I | 1494917 | 127665 | |
| | | | | | | | TR | 1088739 | 159791 | |
| | | | USM | 95 | IgM | no PCR product | | | | |
| | | | SM | 292 | IgG | | | | | |
| | | | | | IgM | | | | | * |
| | | | DN | 96 | IgG | | | | | |
| | | | | | IgM | | | | | |
| | | | PC | 14 | IgG | 3 | I | 407278 | 189715 | |
| | | | | | | | TR | 474419 | 1018 | |
| | | | | | IgM | 1 | I | 60102 | 0 | |

| | | | | | | | | | | |
|---|---|---|---|---|---|---|---|---|---|---|
| | | | | | | | TR | 72897 | 3 | |
| | | PB | Naïve | 200000 | IgM | 16988 | I | 279121 | 60479 | |
| | | | | | | | TR | 351486 | 202860 | |
| | | | USM | 83694 | IgM | 916 | I | 226432 | 118098 | |
| | | | | | | | TR | 336683 | 223244 | |
| | | | SM | 86347 | IgG | 1 | I | 496 | 25 | |
| | | | | | | | TR | 1 | 0 | * |
| | | | | | IgM | 1 | I | 258700 | 59 | |
| | | | | | | | TR | 141 | 84 | |
| | | | DN | 8342 | IgG | 251 | I | 279017 | 176994 | |
| | | | | | | | TR | 264356 | 167374 | |
| | | | | | IgM | 38 | I | 20296 | 2961 | |
| | | | | | | | TR | 25569 | 5624 | |
| | | | PC | 59 | IgG | 3 | I | 314085 | 200898 | |
| | | | | | | | TR | 549128 | 443931 | |
| | | | | | IgM | 4 | I | 225174 | 142723 | |
| | | | | | | | TR | 423936 | 323116 | |
| | T2 | CSF | Naïve | 206 | IgM | 40 | I | 1901177 | 105912 | |
| | | | USM | 218 | IgM | 10 | I | 531781 | 175800 | |
| | | | SM | 648 | IgG | no PCR product | | | | * |
| | | | | | IgM | | | | | |
| | | | DN | 47 | IgG | 5 | I | 462534 | 140602 | |
| | | | | | IgM | 2 | I | 137253 | 62 | |
| | | | PC | 49 | IgG | 3 | I | 632600 | 188073 | |
| | | | | | IgM | 1 | I | 11956 | 16 | |
| | | PB | Naïve | 200000 | IgM | 5779 | I | 339081 | 37038 | |
| | | | USM | 200000 | IgM | 3779 | I | 289896 | 66564 | |
| | | | SM | 200000 | IgG | 2280 | I | 621418 | 173182 | |
| | | | | | IgM | 323 | I | 187316 | 53827 | |
| | | | DN | 49382 | IgG | 2176 | I | 507080 | 123160 | |
| | | | | | IgM | 147 | I | 81149 | 9991 | |
| | | | PC | 3860 | IgG | 66 | I | 375733 | 131626 | |
| | | | | | IgM | 62 | I | 79833 | 30496 | |

On average 2118 (+/- 9953 SD) aligned reads were obtained per cell. Shown are the number of B-cells in each patient's sorted or bulk sample(s) at T1 and T2. Bulk samples contain all B-cells from a given time point sorted into a single sample tube; FACS-sorted B-cell subsets are naïve, USM, SM, DN, or PC. The number of IgG-VH/IgM-VH clusters derived from each sample's IgG-VH and IgM-VH sequencing libraries are shown as well as initial and technical replicate raw sequencing read counts and aligned post-MiXCR read counts. Exp, Experiment: I, initial. TR, technical replicate. FACS, fluorescence-activated cell sorting. *Subsets/Ig isotypes from which no Ig-VH libraries could be obtained. † 5 subsets yielded more Ig-VH clusters than the number of input cells. For these samples, we analyzed the most abundant Ig-VH clusters, such that the number of clusters did not exceed the number of input cells.

**Table S4: CSF Ig-VH cluster persistence rate is similar to PB Ig-VH cluster persistence rate.**

| | **% of T1 Ig-VH clusters (+/- SD)** | **% of T2 Ig-VH clusters (+/- SD)** | **p-value** CSF-persistence vs PB-persistence (T1, T2) |
|---|---|---|---|
| **CSF-persistence rate** patients with persistent CSF Ig-VH clusters | 5.4% (+/- 7.2) | 13.1% (+/- 20.9) | n/a |
| **PB-persistence rate** patients with persistent CSF Ig-VH clusters | 6.4% (+/- 2.9) | 7.9% (+/- 3.2) | p=0.8 and p=0.6 |
| **PB-persistence rate** patients without persistent CSF Ig-VH clusters | 5.6% (+/-3.8) | 4.3% (+/-5.5) | p=1.0 and p=0.5 |

Ig-VH cluster persistence rate is defined as the percent of total Ig-VH clusters from T1 or T2 that are found in both T1 and T2 samples (Persistence rate as % of T1 = # Ig-VH clusters found at both T1 and T2 / Total # Ig-VH clusters at T1). Persistence rate in the CSF is compared to the PB-persistence rate in the five patients with persistent CSF Ig-VH clusters as well as the PB-persistence rate in the five patients without persistent CSF Ig-VH clusters. Unpaired t-tests, $p<0.05$ was considered significant.

**Table S5. FACS antibody sort panels**.

| **Sort panel** | **BV421** | **FITC** | **PerCP-Cy5.5** | **eF710** | **PE-Cy7** | **PE** | **APC** | **APC-Alexa750** |
|---|---|---|---|---|---|---|---|---|
| **3** | CD4 | CD20 | CD19 | --- | --- | CD14 | CD3 | CD8 |
| **6** | IgD | CD20 | CD38 | --- | CD3 | CD138 | CD27 | CD19 |
| **7** | IgD | CD19 | --- | CD5 | --- | CD38 | CD27 | --- |
| **8** | IgD | CXCR5 | CD38 | --- | --- | CD138 | CD27 | CD19 |
| **19** | IgD | CXCR5 | CD38 | --- | CD3 | CD138 | CD27 | CD19 |
| **22** | IgD | CD20 | CD19 | --- | --- | CD3 | CD27 | CD8 |
| **18** | IgD | CXCR5 | CD38 | --- | CD3 | CD138 | CD27 | CD19 |

Panels of fluorescent antibodies used to identify and sort B-cell subpopulations by flow cytometry. IgD Brilliant Violet 421 (Biolegend 11-26c.2a), CD4 Brilliant Violet 421 (Biolegend OKT4), CD20 FITC (Beckman Coulter B9E9), CXCR5 FITC (Biolegend J252D4), CD19 FITC (Biolegend HIB19), CD38 PerCPCy5.5 (BioLegend HIT2), CD19 PC5.5 (Beckman Coulter J3-119), CD5 PerCP-eFluor710 (eBioscience YKIX322.3), CD3 PE-Cy7 (Beckman UCHT1), CD138 PE (Miltenyi 449), CD3 PE (Beckman Coulter UCHT1), CD38 PE (eBioscience 90), CD14 PE (eBioscience 61D3), CD27 APC (eBioscience O323), CD3 APC (Beckman UCHT1), CD19 APC-Alexa750 (Beckman J3-119), CD8 APC-Alexa750 (Beckman B9.11). BV421, brilliant violet 421.

**Table S6. FACS antibody sort panels used on cerebrospinal fluid and peripheral blood**.

| Patient ID | Time point | PB sort panel (1) | PB sort panel (2) | CSF sort panel |
|---|---|---|---|---|
| **1** | T1 | 7 | | 22 |
| | T2 | 18 | | 19 |
| **2** | T1 | 7 | 3 | |
| | T2 | 7 | 3 | 22 |
| **3** | T1 | 7 | | |
| | T2 | 18 | | 19 |
| **4** | T1 | 7 | 3 | 22 |
| | T2 | 7 | 3 | 6 |
| **5** | T1 | 7 | 3 | |
| | T2 | 7 | 3 | |
| **6** | T1 | 7 | 3 | 8 |
| | T2 | 18 | | 19 |
| **7** | T1 | 7 | 3 | |
| | T2 | 7 | 3 | |
| **8** | T1 | 7 | 3 | 8 |
| | T2 | 18 | 3 | 19 |
| **9** | T1 | | | |
| | T2 | 18 | | 19 |
| **10** | T1 | 7 | 3 | 8 |
| | T2 | 18 | | 19 |

Panels of fluorescent flow cytometry antibodies that were used on each sample at each time point. See Table S5 for details of each sort panel.
